# Nuclear pores as versatile reference standards for quantitative superresolution microscopy

**DOI:** 10.1101/582668

**Authors:** Jervis Vermal Thevathasan, Maurice Kahnwald, Konstanty Cieśliński, Philipp Hoess, Sudheer Kumar Peneti, Manuel Reitberger, Daniel Heid, Krishna Chaitanya Kasuba, Sarah Janice Hoerner, Yiming Li, Yu-Le Wu, Markus Mund, Ulf Matti, Pedro Matos Pereira, Ricardo Henriques, Bianca Nijmeijer, Moritz Kueblbeck, Vilma Jimenez Sabinina, Jan Ellenberg, Jonas Ries

**Author notes:** Current affiliation: Newcastle University (Sudheer Kumar Peneti). Current affiliation: DKFZ, Heidelberg. Current affiliation: Department for Applied Tumor Biology, Heidelberg University Hospital (Daniel Heid). Current affiliation: D-BSSE, ETH Zürich. Current affiliation: Institute of Molecular and Cell Biology, Mannheim University of Applied Science and Interdisciplinary Center for Neuroscience, Heidelberg University (Sarah Janice Hoerner). Current affiliation: Department of Biochemistry, University of Geneva, Science 2, Quai Ernest-Ansermet 30, CH-1205 Genève, Switzerland. these authors contributed equally (Jervis Vermal Thevathasan, Maurice Kahnwald).

## Abstract

Quantitative fluorescence and superresolution microscopy are often limited by insufficient data quality or artifacts. In this context, it is essential to have biologically relevant control samples to benchmark and optimize the quality of microscopes, labels and imaging conditions.

Here we exploit the stereotypic arrangement of proteins in the nuclear pore complex as in situ reference structures to characterize the performance of a variety of microscopy modalities. We created four genome edited cell lines in which we endogenously labeled the nucleoporin Nup96 with mEGFP, SNAP-tag or HaloTag or the photoconvertible fluorescent protein mMaple. We demonstrate their use a) as 3D resolution standards for calibration and quality control, b) to quantify absolute labeling efficiencies and c) as precise reference standards for molecular counting.

These cell lines will enable the broad community to assess the quality of their microscopes and labels, and to perform quantitative, absolute measurements.

## Introduction

During the last decade, superresolution microscopy, specifically single-molecule localization microscopy (SMLM, also known as (f)PALM^1,2^ or STORM^3^), has pushed the resolution of optical microscopy towards the nanometer scale and has provided structural insights into cell biological questions^4–6^. To successfully use SMLM, many factors have to be optimized: microscope optics, settings and stability; imaging conditions such as buffer composition; characteristics of fluorophores and labeling technologies; sample preparation protocols; and analysis software. Although these factors are crucial to obtain high quality SMLM images, to date a common practice of quality control that ensures comparable quality of superresolution data between different labs, or even within a lab is missing. Suboptimal performance in SMLM is therefore not readily detected, which severely limits biological interpretation and discovery, and wastes resources. There are algorithms that evaluate the quality in some microscopy modalities^7,8^, but without knowledge about the underlying structure being imaged they are limited in their quality assessment.

This limitation can be overcome by suitable reference samples, which allow a quantitative characterization and systematic optimization of a superresolution microscopy workflow. This reference-based approach has been used successfully to benchmark various superresolution analysis software packages, where simulated synthetic images served as reference standards^9^. In the past, both artificial and cellular structures have been used as physical references. While artificial structures such as DNA origami^10,11^ allow positioning fluorophores at precise three-dimensional positions, they are limited in the choice of labels and are intrinsically different from intra-cellular biological structures. Cellular reference structures have included histones^2,12^, mitochondria^13^, and, most commonly, microtubules^14^. For instance, a widely used approach to approximate the experimental resolution relies on evaluating cross-sectional profiles of manually selected microtubules, which however is prone to artifacts and cherry-picking^15^. One major limitation of these references is their abundance of epitopes. This results in acceptable images even for labeling efficiencies below 1%. Therefore, these references cannot be used to characterize and optimize labeling efficiencies, a major factor determining image quality in SMLM.

An ideal superresolution reference structure should fulfill the following requirements: its fluorescent labels are precisely positioned in 3D at distances that are resolvable by the technique of choice; it contains a defined number of fluorophores to allow quantifying labeling efficiencies; it is compatible with common dyes and labeling approaches in order to meaningfully resemble the actual intra-cellular measurements; it is present in many copies in the cell for statistical accuracy; and finally it is simple and reproducible to prepare.

The nuclear pore complex (NPC) fulfills all of these requirements and is thus particularly well-suited as a quantitative reference structure. The NPC is a large protein complex in eukaryotes that comprises ~30 different proteins, so-called nucleoporins, and ensures selective macromolecular transfer across the nuclear membrane. The human NPC has been extensively studied by electron microscopy^16^, yielding a high resolution structural map of most nucleoporins. Several superresolution microscopy studies have provided additional insight into nanoscale arrangement and abundance of numerous nucleoporins^6,17,18^.

Here, we generated cell lines where the nucleoporin Nup96 is endogenously tagged with commonly used labels. We demonstrate that imaging these cell lines yields excellent reference data to control the experimental parameters that are essential for quantitative superresolution microscopy. We show in detail how they can be used to a) quantify microscope performance, resolution and spatial calibration, b) measure absolute labeling efficiencies, c) optimize imaging conditions, and d) count the number of proteins within a complex.

## Results

### Generation of Nup96 cell lines

Among the ~30 proteins in the NPC, we identified that Nup96 is ideally suited to serve as a reference protein (**Fig. 1a-e**). Nup96 is present in 32 copies per NPC where it forms a cytoplasmic and a nucleoplasmic ring, each consisting of 16 Nup96 copies. Each ring has 8 corners that contain two Nup96, 12 nm apart^16^. The nucleoplasmic and cytoplasmic rings of Nup96 are almost in register, thus in a top view the characteristic eightfold symmetry of the NPC is clearly visible (**Fig. 1e**).

**Figure 1:**
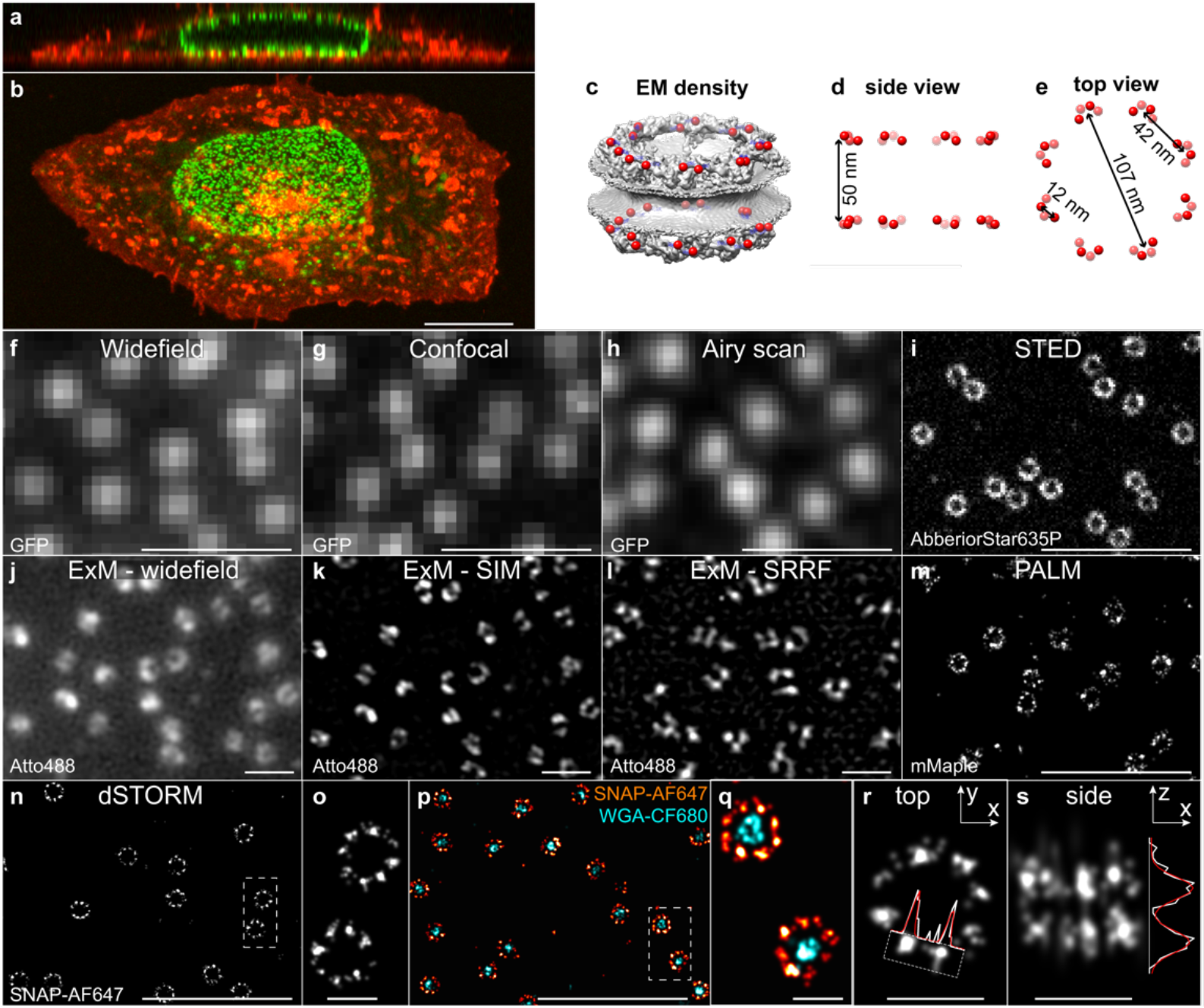
Nup96 cell lines. (**a**) confocal x-z and (**b**) x-y image of the Nup96-GFP cell line. green: Nup96-GFP, red: membranes (DiD). (**c**) EM density of the nuclear pore complex^16^ with C-termini of Nup96 indicated in red. (**d**) Side view and (**e**) top view schematic. (**f**) widefield, (**g**) confocal and (**h**) airy scan images of Nup96-GFP. (**i**) STED image of Nup96-GFP labeled with an AberriorStar635P-coupled anti-GFP nanobody. Resolution estimates based on Fourier power spectra for **f-i** can be found in **Supplementary Figure 2a.** (**j**) Widefield expansion microscopy image of Nup96-GFP labeled with an Atto488-coupled anti-GFP nanobody. (**k**) as before, but imaged using structured illumination. Estimates of the expansion factor based on the analysis of the ring diameters can be found in **Supplementary Figure 2c.** (**l**) as before, but imaged using SRRF. (**m**) SMLM image of Nup96-mMaple, (**n, o**) SMLM of Nup96-SNAP labeled with BG-AF647. (**p, q**) Dual-color SMLM image of Nup96-SNAP labeled with BG-AF647 (red) and WGA-CF680 (cyan). (**r, s**) Corners of the NPC can be used as a resolution target in x,y (**r**) and z (**s**). Resolution estimates based on Fourier Ring Correlation for **m-q** can be found in **Supplementary Figure 2b**. Scale bars 10 μm (**b**), 1 μm (**f-n,p**) and 100 nm (**o,q,r,s**).

We generated four homozygous CRISPR knock-in cell lines^19^ in which Nup96 is endogenously labeled with one of four commonly used labels: mEGFP (subsequently referred to as GFP), the photoconvertible fluorescent protein mMaple^20^, and the enzymatic labels SNAP-tag^21^ and HaloTag^22^. Homozygosity of all cell lines was validated by Sanger sequencing, Southern and Western blot (**Supplementary Figure 1**). We chose U2OS cells as they are well suited for microscopy because of their flatness. Their lower nuclear envelope is close to the coverslip (**Fig. 1a**), thus hundreds of nuclear pores can be in focus at the same time, and they can be imaged using widefield, confocal or total internal reflection (TIRF, **Supplementary Figure 3**) excitation.

### Resolution and quality control

Our reference cell lines position fluorophores at distinct distances ranging from 10 nm to 100 nm, and are therefore suitable resolution standards for many microscopes. We first imaged them with 9 different microscopy approaches widely used in biological imaging (**Fig. 1f-n**). We obtained images with excellent signal to noise ratio for all imaging modalities, highlighting the differences in resolving power of each microscope. For diffraction-limited techniques, NPCs act as sub-diffraction structures of defined brightness. Individual NPCs are resolved in widefield microscopy (**Fig. 1f**), and, with improved contrast in confocal microscopy (**Fig. 1g**). Airy-scan microscopy (**Fig. 1h**) leads to a visible, but moderate improvement in resolution (**Supplementary Figure 2a**). Stimulated emission depletion (STED^23^) microscopy reaches a higher resolution and clearly resolves the ring-like arrangement of Nup96 (**Fig. 1i**). These rings are also apparent in expansion microscopy^24^ with widefield (**Fig. 1j**), structured illumination^25^ (**Fig. 1k**) and SRRF^26^ (**Fig. 1l**) readout. In expansion microscopy the local expansion factor is of key importance for quantitative analyses, but in general it is difficult to calibrate. In our reference cell lines, however, it can be directly inferred from the size of the rings^27^ (Methods). We found that the local expansion factor of 3.2 was different from the global expansion factor of 4.5 determined from the size change of the sample upon expansion, indicating heterogeneities in expansion (**Supplementary Figure 2c**).

SMLM using mMaple (PALM^2^ approach) shows clear rings of the NPC and starts resolving the eight corners (**Fig. 1m**), even when imaged in living cells (**Supplementary Figure 4**). The highest resolution is reached using organic dyes (*d*STORM approach^28^) (**Fig. 1n,o**), where the eight corners are very well resolved (**Fig. 1r**), also in dual-color superresolution images (**Fig. 1p,q**). The visual impression of increasing lateral resolution was confirmed in the Fourier spectrum^29^ (**Supplementary Figure 2a**) and by Fourier Ring Correlation^30^ (**Supplementary Figure 2b**).

Our cell lines are also optimally suited to quantify the axial resolution of 3D superresolution imaging. If the z-resolution is high enough, the two rings of the NPC can be resolved in an axial profile, where the standard deviation of each peak is an upper limit for the experimental localization precision (**Fig. 1s**, **Fig. 2h,j**).

**Figure 2:**
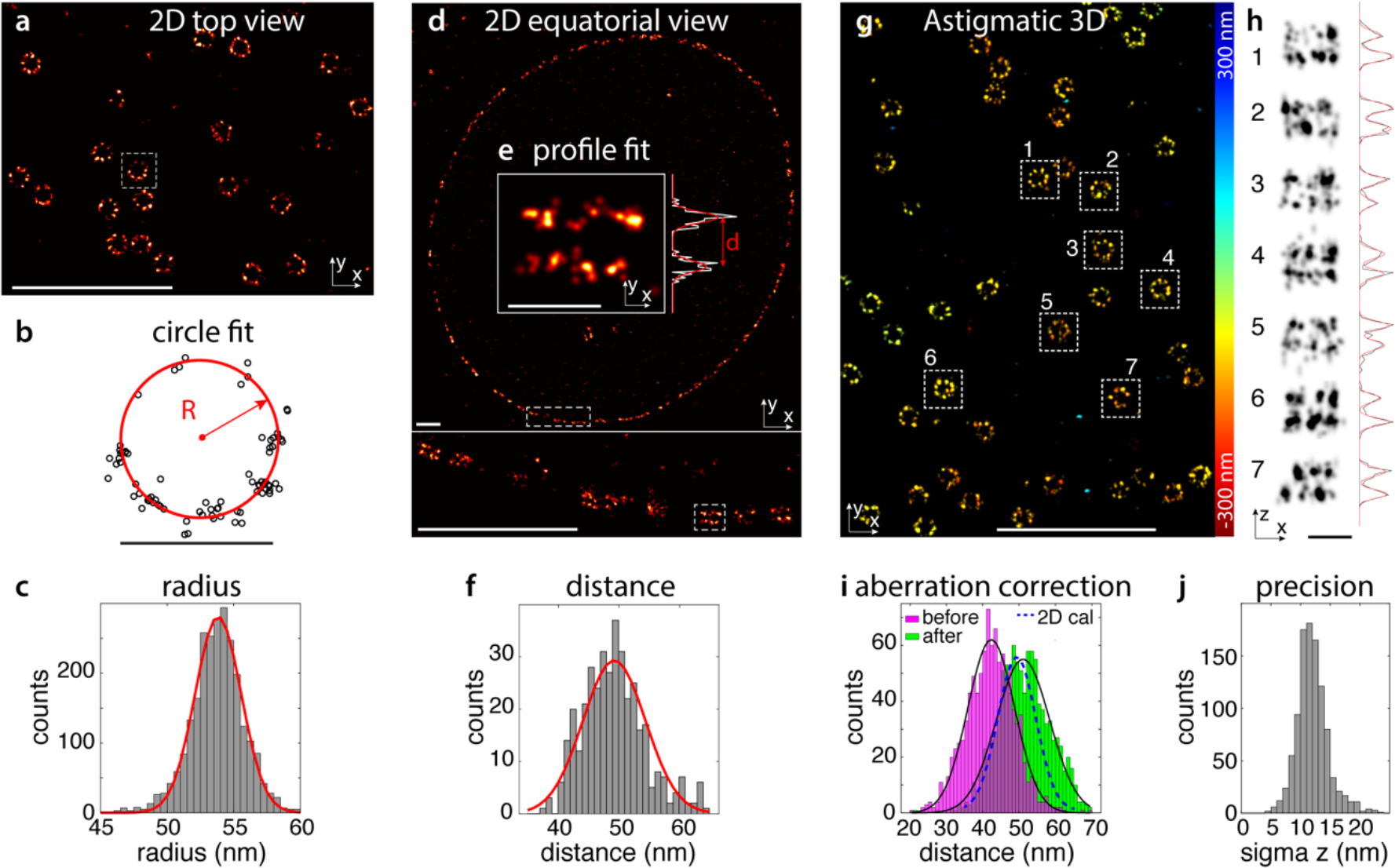
Nuclear pores as calibration reference standards. (**a-h**) **experimental characterization of Nup-96 positions in the NPC.** (**a**) SMLM image of lower nuclear envelope, (**b**) circle fit of a single NPC, (**c**) histogram of fitted radii (R = 53.7 nm (mean) ± 2.1 nm (SD), N = 2536 NPCs). (**d**) equatorial SMLM image of Nup96, (**e**) a single NPC in a side view. A fit with a double Gaussian returns the ring-distance d and the standard deviation of each ring. (**f**) histogram of separation between rings (d = 49.3 nm ± 5.2 nm, N = 379 NPCs). (**g**) 3D SMLM image of lower nuclear envelope. The localizations are color-coded according to their z-position. (**h**) x-z reconstructions with z-profiles as indicated. (**i**) **NPCs as calibration reference standard for astigmatic 3D SMLM.** Histogram of ring-distances before correction (magenta, d = 42.1 ± 1.1 nm, N = 8 cells) and after correcting for depth-induced calibration errors (green, d = 49.8 ± 1.9 nm, N = 8 cells). (**j**) Standard deviation of z-profiles from double Gaussian fit result in an upper bound for the experimental localization precision in z of 13.3 ± 1.0 nm (N = 8 cells). Scale bars 1 μm (**a,d,g**), 100 nm (**b,e,h**). All data on Nup96-SNAP-AF647.

### Microscope calibration

Nup96 is precisely arranged within the NPC. Thus, the calibration of a superresolution microscope can be verified by comparing measured distances between individual Nup96 clusters to the known distances within the complex. As each nucleus contains hundreds to thousands of NPCs, automated analysis of a single image yields a high precision estimate.

We first measured all relevant distances within the NPC using an SMLM microscope with a precisely calibrated pixel size (Methods). The average radius of the NPC was R = 53.7 ± 2.1 nm (**Fig. 2a-c**, all values in this manuscript are mean ± standard deviation (SD) unless otherwise noted). We then acquired side-view images of the NPCs by focusing on the midplane of the nucleus, and determined that the cytoplasmic and nucleoplasmic rings were 49.3 ± 5.2 nm apart (**Fig. 2d-f**, Methods). Using 3D SMLM (**Fig. 2g,h**), we determined an azimuthal shift of 8.8° ± 0.6° between both rings using angular cross-correlation (Methods). Our measurements agree well with the EM structure^16^, with minor deviations, probably because the C-terminus of Nup96 tagged here was not modeled in the EM structure.

With the dimensions of Nup96 in the NPC now precisely calibrated, our cell lines can be used to verify the pixel size calibration of any superresolution microscope by analyzing the diameter of many rings and comparing the average value to the value as stated above. In addition to the pixel size, our cell lines can be used to verify the axial calibration in 3D SMLM. This is typically challenging, as aberrations or imperfect calibration of the PSF model can lead to systematic and depth-dependent localization errors, especially when using oil objectives^31^. Moreover, the refractive index difference between oil and the aqueous sample leads to an image compressed in z. This can be corrected by applying a refractive index mismatch factor (RIMF)^14^, which however is usually not precisely known and difficult to calibrate. Our tagged Nup96 cell lines overcome this limitation and enable a quantitative analysis of the z-calibration and the RIMF. For this, we checked the z-calibration of our astigmatic SMLM microscope by measuring the distance between the two rings in thousands of NPCs in 3D. The average distance was d = 42.2 ± 1.2 nm (**Fig. 2i**, based on a RIMF of 0.8), and thus smaller than the true value of 49.3 nm. Furthermore, the distance between the rings was correlated to the distance between NPC and coverslip (**Supplementary Figure 5a**), indicating aberrations. After correcting for these aberration-induced fitting errors with a method we recently developed^31^, the z-dependence was reduced (**Supplementary Figure 5b**) and the average corrected distance is d = 49.8 ± 1.9 nm. Based on these results we could calibrate the RIMF to be 0.79.

### Effective labeling efficiencies

Besides the localization precision, the information content of any SMLM image most importantly depends on how densely the structure of interest is decorated with fluorophores. This can be described by the *effective* labeling efficiency (ELE), which represents the fraction of target proteins that carry a fluorophore that is detected as a useable localization (i.e. brightness above background, fitted with acceptable confidence).

Generally, the ELE is hard to quantify, as the number of proteins in the target structure is usually unknown, and a higher ELE is difficult to distinguish from increased blinking and re-activation of the fluorophores. In the case of photoactivatable proteins, protein and chromophore maturation have to be accounted for. The maturation efficiency of photoactivatable proteins has been estimated using receptors on the cell surface^32^ or by mathematical modeling of fluorophore photo physics.^33^ Intricate experiments have been developed to determine binding efficiencies of anti-GFP antibodies^34^. Altogether, despite its critical importance, a simple and robust approach to measure the ELE of common labeling strategies inside cells is still missing. This has severely limited our ability to systematically optimize image quality in SMLM. Furthermore, the ELE of many new fluorophores or labeling strategies are not well characterized and their performance is often not as ideal as in proof-of-concept experiments on abundant proteins.

Our Nup96 cell lines provide the unique opportunity to directly measure the absolute ELEs of GFP-binders and SNAP-tag or HaloTag ligands with a simple assay. When the ELE is low, NPCs appear as incomplete rings with missing corners. Thus, by counting the number of corners for many NPCs, we can infer the absolute ELE from a statistical analysis with a probabilistic model. Here, we developed a straightforward workflow to determine the number of corners in hundreds of NPCs based on automatic segmentation, registration to a template and counting of areas that contain at least one localization (**Fig. 3**, Methods). The variability among different cells and biological replicates was typically smaller than 10% (SD, **Fig 3i**). Using simulations (**Supplementary Figure 6**), we showed that this approach is robust over a large range of ELEs, localization precisions and number of re-activations.

**Figure 3:**
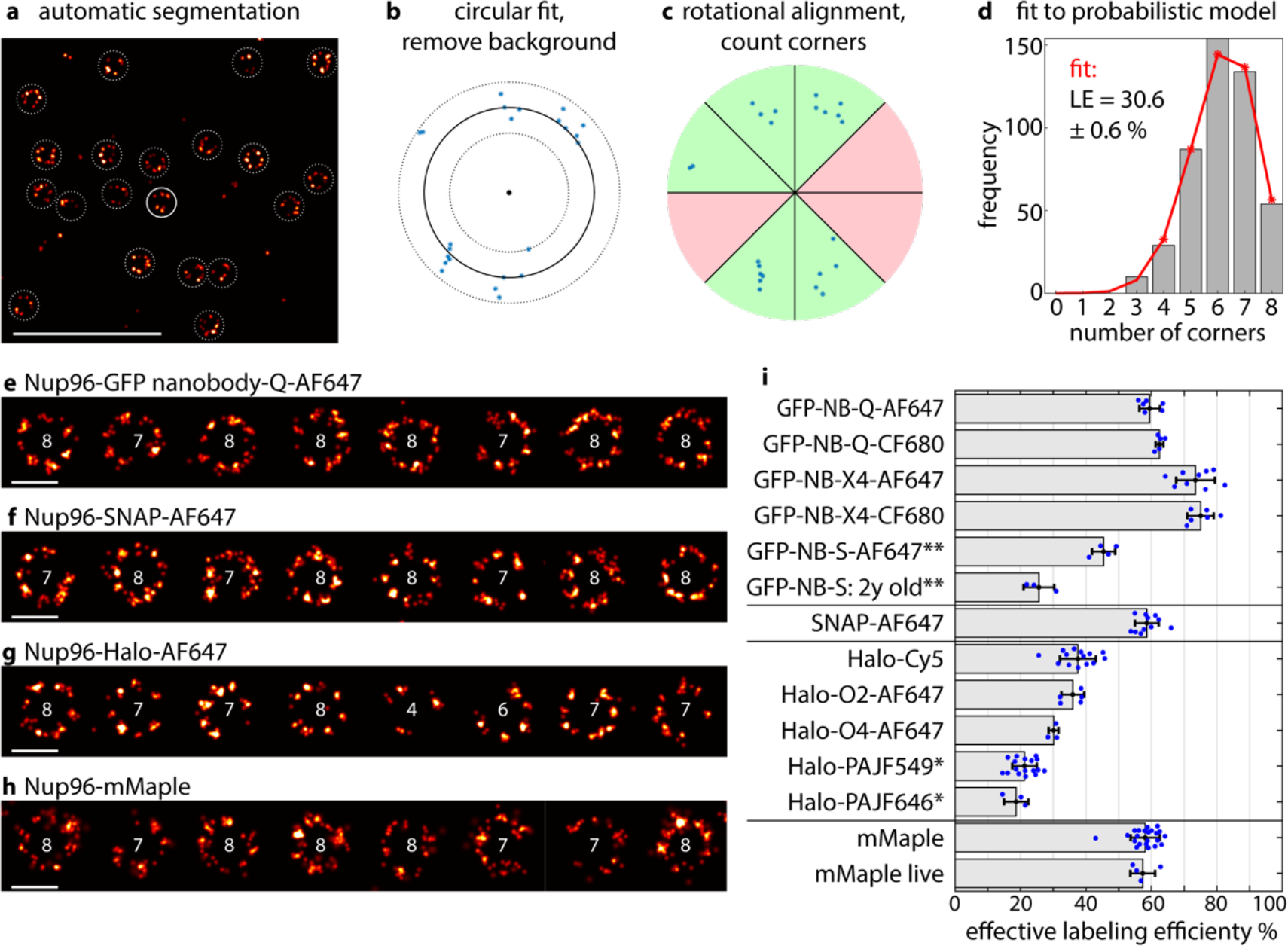
Effective labeling efficiencies. **(a-d) workflow**. (**a**) All NPCs in a cell are automatically segmented. (**b**) We fit a circle to the localizations and reject localizations outside a ring as background localizations (**d**) We rotate the localizations to optimally fit an eightfold-symmetric template and count the number of slices that contain at least one localization. (**d**) We fit the histogram of the number of corners with a probabilistic model to directly obtain the absolute ELE. The statistical error is estimated by bootstrapping. (**e-h**) **gallery of NPCs.** (**e**) **Nup96-GFP** labeled with an anti-GFP nanobody coupled to AF647. (**f**) **Nup96-SNAP** labeled with BG-AF647. (**g**) **Nup96-Halo** labeled with chloroalkane-AF647. (**h**) **Nup96-mMaple.** The numbers indicate the numbers of visible corners the algorithm detected. (**i**) Effective labeling efficiencies for various cell lines and ligands. Bars denote the mean, error bars the standard deviation and individual data points measurements of a single cell. Scale bars 1 μm (**a**) and 100 nm (**e-h**). *labeled in live cells, imaged after fixation. **measured on Nup107-GFP.

Using this analysis pipeline, we systematically compared the effective labeling efficiencies of different anti-GFP nanobodies^35^, and SNAP-tag and HaloTag ligands with different organic dyes (**Fig. 3i**, **Supplementary Table 1**). We observed the highest ELEs of ~74% using commercial anti-GFP nanobodies that contain a mixture of two different nanobodies. Other monoclonal commercial anti-GFP nanobodies achieved ~60%, while anti-GFP nanobodies that we generated in the lab showed a lower ELE of 45%, which was further reduced to 25% after 2 years of storage in the fridge.

For Nup96-SNAP labeled with BG-AF647 we achieved an ELE of 60%. Due to the covalent nature of binding of BG to SNAP, labeled and fixed samples were stable over years (demonstrated on an analogous Nup107-SNAP cell line, **Supplementary Figure 7**) with only minor loss in ELE. This ability to prepare and store standard samples greatly facilitates their prolonged regular usage.

Using HaloTag, we achieved substantially lower ELEs of 19% – 38% with five different fluorescent ligands, and found that choice of fluorophore and linker length strongly influenced the efficiency. While the photo-activatable ligands PA-JF549 and PA-JF646^36^ showed no specific labeling in fixed cells, they could be used for live-cell labeling with ELEs of 21.1 ± 3.7 % for PA-JF549 and of 18.7 ± 3.0 % for PA-JF646. In addition to the ELE, our approach reveals how often a single fluorophore is detected per experiment (Methods). For instance, while a single AF647 dye is localized on average 4.4 ± 0.7 times, PA-JF549 produces on average 1.3 ± 0.1 localizations and thus shows little blinking. This is well-suited to investigate protein clustering as it reduces false positives caused by re-activation of fluorophores.

Using simulations (**Supplementary Figure 6**) we showed that our approach of quantifying the ELE works even when the individual corners are not always clearly discernible. Thus, we extended our analysis to the photoconvertible fluorescent protein mMaple. We found an ELE of 58% ± 4 %, indicating that even though 100% of all Nup96 are fused to mMaple, about 40% of them are not detected as a localization. This is likely due to improper folding, insufficient brightness or incomplete photoconversion, in line with previous reports^32,33^.

Taken together, this assay provides an easy way for any lab using SMLM to monitor the ELEs of their labeling reagents, including nanobodies, SNAP-tag and HaloTag ligands, thus avoiding the use of sub-optimal labels.

### Imaging conditions

In any SMLM measurement, numerous factors influence the quality of the final image, including imaging buffers, laser power densities, exposure times, choice of filters, and settings in the analysis software. To find optimal conditions, these factors are typically varied while optimizing for various read-outs for quality, including brightness of single fluorophores, low background, on-times, duty cycle, localization precision, ELE, number of re-activations, imaging speed or stability of imaging buffers. Such an optimization requires a robust standard sample with small variability to allow detection of subtle changes.

Our Nup96 cell lines allow for robust read-out of these quality-related parameters and we demonstrated their use by comparing several imaging conditions (**Table 1**). We confirmed that AF647 is hardly affected by D_2_O in solution^37^, and that, compared to the MEA buffer, it shows increased brightness and number of localizations per fluorophore in BME^38^, and reduced brightness and number of localizations per fluorophore in sulfite buffer^39^. Interestingly, we found that a high ELE correlates with a large number of localizations per fluorophore, a possible explanation for this is bleaching during the first switching-off cycle.

**Table 1:** Imaging conditions. Effective labeling efficiency, mean photons per localization and mean localizations per fluorophore for Nup96-SNAP-AF647 and Nup96-mMaple in commonly used imaging buffers. All values are mean ± SD (N ≥ 4 cells).

| Sample | Buffers | Effective LE % | Photons per localization | Localizations per fluorophore |
| --- | --- | --- | --- | --- |
| SNAP-AF647 | 35 mM MEA + GLOX | $60 \pm 3$ | $10674 \pm 833$ | $4.4 \pm 0.7$ |
| | 35 mM MEA + GLOX in D<sub>2</sub>O | $56 \pm 3$ | $9986 \pm 365$ | $3.6 \pm 0.4$ |
| | 143 mM BME + GLOX | $63 \pm 4$ | $12815 \pm 824$ | $5.6 \pm 0.5$ |
| | 35 mM MEA + 50 mM sodium sulfite | $39 \pm 2$ | $6984 \pm 611$ | $1.5 \pm 0.1$ |
| mMaple fixed | 50 mM Tris in D<sub>2</sub>O | $58 \pm 4$ | $2159 \pm 259$ | $2.7 \pm 0.2$ |
| mMaple live | 50 mM Tris in D<sub>2</sub>O | $57 \pm 3$ | $1877 \pm 50$ | $2.8 \pm 0.1$ |

Finally, we found that fixation with PFA did not have a substantial effect on mMaple photophysics or ELE.

### Counting of proteins

The stoichiometry of a multi-protein assembly, i.e. the absolute number of subunits, is an essential parameter for studying multiple aspects of its function. Converting gray values from a fluorescence microscopy image to absolute protein numbers requires careful calibration of the microscope. The Nup96-GFP cell line is well suited for this task, as we can resolve the majority of nuclear pores even in diffraction-limited microscopy (**Fig. 1f,g**, **Fig. 4a**), and thus can calibrate precisely how bright 32 GFP-labeled proteins are. We can then use this calibration to determine the unknown abundance of a different GFP-labeled protein. For validation, we chose Nup107, which is another nucleoporin that is present in 32 copies per NPC^40^. We used a simple brightness analysis in which we evaluated the intensity of the brightest pixel of a local intensity maximum as a measure for the brightness of the NPC and found similar average values for the Nup96-GFP and Nup107-GFP in a HEK cell line^41^ (**Fig. 4a-d**) with relative errors below 5% SD.

**Figure 4:**
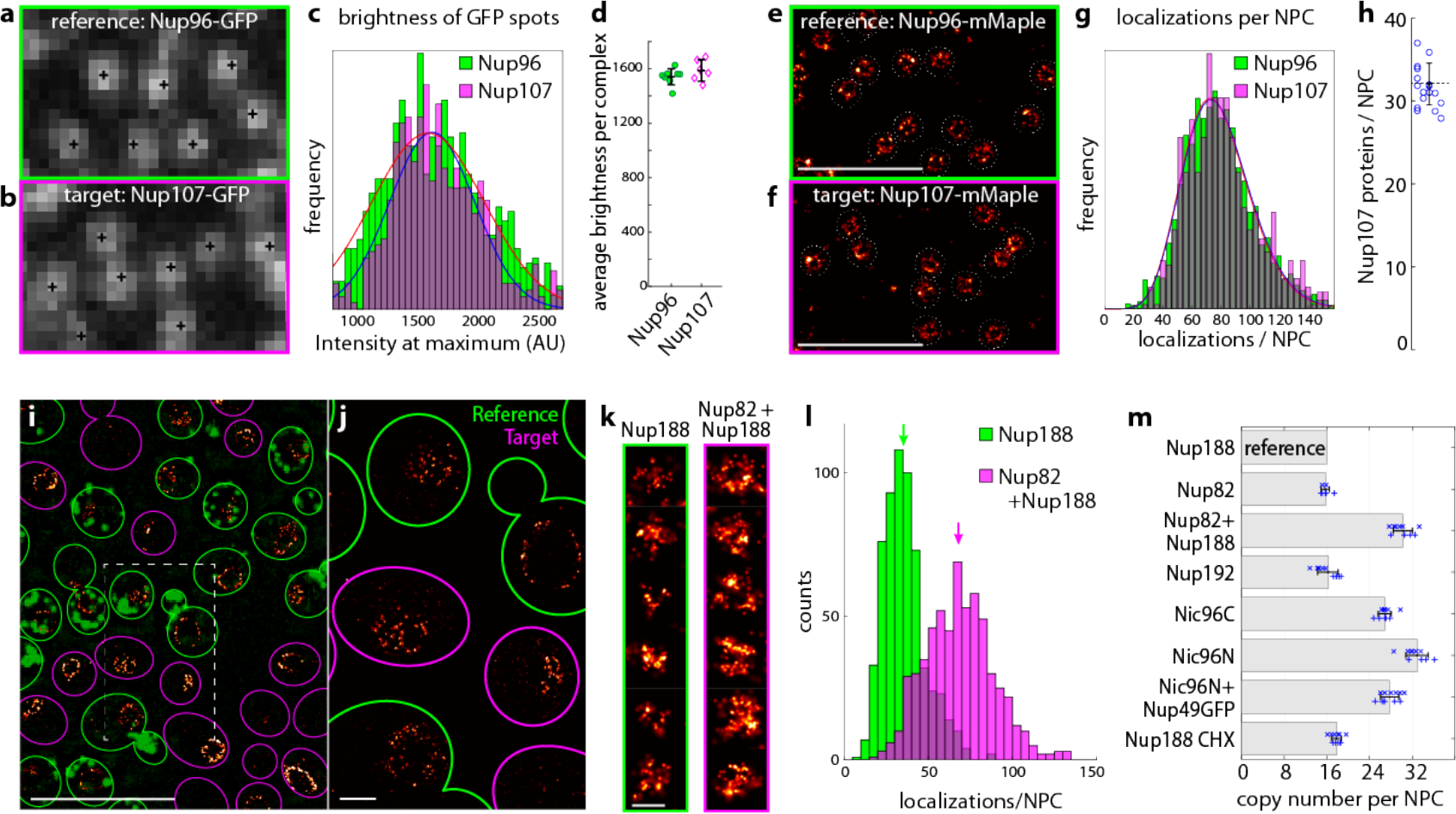
Counting of protein copy numbers in complexes. (**a-d**) **Counting in diffraction limited microscopy.** (**a**) confocal image of the reference protein Nup96-GFP with the majority of nuclear pores resolved. (**b**) confocal image of the target protein Nup107-GFP imaged with the same microscope settings. (**c**) histograms of intensities of local maxima (see Methods) for the reference and target structures together with Gaussian fit to determine the mean intensity values. (**d**) mean intensity values for several reference and target cells. These values show a small variation and are similar for reference (〈*I*_ref_〉 = 1541 (mean) ± 59 (SD) ADU, N = 8 cells) and target complex (〈*I*_tar_〉 = 1593 ± 79 ADU, N = 6 cells). (**e-h**) **Counting with SMLM.** (**e**) reconstructed superresolution for reference cell line Nup96-mMaple and (**f**) for target cell line Nup107-mMaple. NPC structures are automatically segmented to determine the numbers of localizations per NPC. (**g**) Histogram of number of localizations per NPC for reference and target. The number of Nup107-mMaple proteins per NPC is calculated from the average relative number of localizations. (**h**) The stoichiometry of Nup107 in the NPC (*N*_Nup107_ = 32.2 ± 2.5, mean ± SD, N = 17 cells) shows a high accuracy and low statistical errors of this counting approach. (**i-m**) **Counting in yeast.** (**i**) Mixture of Nup188-mMaple+Abp1-GFP reference cell lines with Nup82-mMaple+Nup188-mMaple target cell lines, which can be distinguished by the GFP signal. (**j**) superresolution reconstruction and (**k**) individual nuclear pores. (**l**) histograms of the number of localizations per nuclear pore, arrows indicate the mean. (**m**) copy number of several yeast nucleoporins per NPC, determined using Nup188 as a reference.

Extracting absolute molecular numbers in superresolution is especially beneficial to unravel stoichiometries of small and dense multi-protein assemblies. However, relating the number of localizations to the number of proteins is not trivial in SMLM. Incomplete labeling leads to undercounting, while repeated re-activation of the same fluorophore leads to overcounting. Previous approaches have relied on calibrating blinking and other photophysical properties of fluorophores^42–44^, which cannot account for long-lived dark states and incomplete labeling or maturation. Furthermore, a variety of counting references have been developed including self-assembling oligomers^45^, DNA-structures^34^, receptors^32^, or a combination of monomers, dimers and trimers of mEos2^33,46^. While this is a powerful approach, a major limitation is the need for faithful segmentation, which is not trivial and often strongly dependent on algorithmic parameters. Background localizations or incompletely assembled or labeled reference structures lead to an underestimation of the reference brightness, whereas fusion of double-structures or a cutoff during segmentation and thus loss of small structures leads to an overestimation. Moreover, the detection probability of a fluorophore depends on its z-position, which renders cytoplasmic reference structures less accurate.

Our Nup96 cell lines overcome many of these limitations. NPCs have a characteristic shape and large size, never overlap and are thus easy to segment. They are abundant for improved statistics and are at defined z-positions. Thus, our Nup96 cell lines promise to be robust and precise counting reference structures. To validate the use of Nup96-mMaple as a counting reference standard, we generated a stable HEK293T cell line using the Flp-in™ T-REx system in which Nup107-mMaple was overexpressed, while endogenous Nup107 was knocked down (**Supplementary Figure 8**). As expected, we found 32.2 ± 2.5 Nup107 molecules per NPC (**Fig. 4e-h**), highlighting the consistency of this counting approach.

For accurate counting, all target proteins need to carry a label. Such homozygous endogenous protein tagging is laborious and time-consuming in mammalian cells. In yeast, on the contrary, homologous recombination allows for fast and efficient endogenous labeling. Thus, to enable straightforward counting of multiple proteins, we extended our counting references to *S. cerevisiae*. We chose the nucleoporin Nup188 as the reference standard, which is present in 16 copies per NPC^47,48^. For this, we endogenously tagged Nup188 with mMaple in yeast cells that additionally express a GFP-marker at endocytic sites (Abp1-GFP) for easy identification. This allowed us to simultaneously image reference and target cells in the same field of view, which further reduces potential bias that could arise from varying imaging parameters (**Fig. 4i,j**). We validated this approach by counting Nup82, known to be present also in 16 copies per NPC^47,48^, and by counting Nup82 and Nup188 molecules together within a strain where both were tagged with mMaple (**Fig. 4k,l**). The measured copy numbers of 15.7 ± 0.8 and 30.2 ± 1.8 agree well with their expected values of 16 and 32, respectively.

We went on to determine the copy number of additional nucleoporins Nup192 and Nic96 (**Fig. 4m**). Nup192 was found in 16.2 ± 1.9 copies per NPC, agreeing with previous reports^47,48^. Intriguingly, for Nic96, we found 26.8 ± 1.2 copies when Nic96 was tagged at the C-terminus, contradicting previous reports that found 32 copies of Nic96^47^. It was recently proposed that C-terminal tagging impedes Nic96 function^48^, and indeed we measured 32.9 ± 2.1 copies of Nic96 that was endogenously tagged at its N-terminus. When we introduced an additional GFP tag at the C-terminus of Nup49, which interacts with the C-terminus of Nic96, we again measured only 27.8 ± 1.7 copies of Nic96 even when it was tagged at its N-terminus. Our findings demonstrate the reproducibility of our method, and emphasize the risk of tagging artifacts. Careful quantification of proteins with our counting approach offers an experimental avenue to systematically control for them.

As yeast cells have a duplication time of ~ 2 h, maturation times of fluorescent proteins must be considered. Assuming a maturation time of 48 minutes for mMaple^49^ would result in 28% of unmatured mMaple in the steady-state (Methods). To experimentally test the influence of maturation on our copy number measurements, we stopped global protein synthesis by cycloheximide (CHX), reasoning that mMaple synthesized before the treatment should mature to completion. Indeed, when compared to untreated cells, we found a slight increase in localizations by 11 ± 6%. This is less than estimated above, hinting either to a delay in incorporation of Nup188 into the NPC, to a faster maturation time than previously estimated, or to a maturation of mMaple after fixation. Generally, we recommend using this or a related approach to stop protein synthesis whenever the lifetime of the target protein is short or unknown.

## Discussion

By homozygously labeling Nup96 with four common tags, we generated reference standards for a variety of important applications in microscopy. Shared together with the software to perform all analyses, the cell lines will enable many labs to regularly benchmark their microscopes in terms of resolution and calibration, to optimize imaging conditions with high sensitivity, to determine effective labeling efficiencies of their dyes and labels and to count protein copy numbers in complexes.

The assays presented here are robust and reproducible due to the stereotypic architecture of the NPC. Interestingly, we observed some biological variation in the dimension of the NPC structure, in line with previous reports by electron microscopy^50^ (**Fig. 2**). Thus, a statistical analysis of many NPCs is needed for accurate parameter estimates. This heterogeneity might be interesting with respect to nuclear pore biology, and the data accompanying this manuscript could be the basis for such analysis. Although apparent, this structural variability is still smaller than that of 3D DNA origami standard samples^11,51^. Depending on the labeling protocol, we also observed a cell-to-cell variability of the ELE with a subset of cells showing reduced labeling, stressing the need for replicates and optimal sample preparation. Finally, artifacts (e.g. by drift or over-activation) or insufficient localization precision impede accurate determination of ELEs.

It is curious that intracellular labeling with SNAP-tag or HaloTag did not result in complete labeling, although *in vitro* both can be conjugated to completion^21,22,52^. Also, with anti-GFP nanobodies labeling was not complete. This can be due to incomplete folding of the enzymatic tags, inhibition of the tags by fixatives or intracellular components, or by incomplete activation and detection of the fluorophores, imperfect ligands, or bleaching during the initial off-switching step in SMLM, and warrants further investigation and optimization. Incomplete intracellular labeling for SNAP-tag and HaloTag was reported previously^53^, with the choice of dye strongly affecting the ELE. Also the incomplete maturation of photoconvertible proteins has been reported before^32^ and should be a target for further optimization.

As counting reference standards, NPCs are advantageous to small and globular structures due to the ease of segmentation and their defined z-positions. However, some fundamental limitations still apply to any reference-based approach. Counting can only be accurate if all target proteins carry a label, requiring complete replacement of endogenous proteins. Also, if reference and target protein have different turnover rates in the cell, the fraction of matured labels might be different. In this case it might be necessary to globally arrest protein synthesis for a limited time to allow all labels to mature. Finally, counting with fluorescent proteins, as we have used here, is preferable to counting with external labels, as the latter will suffer from different accessibilities of the tags between reference and target.

The Nup96 cell lines optimally complement the current standard sample for SMLM, i.e. immunolabeled microtubules, as they have a defined stoichiometry and 3D arrangement of the fluorophores and are compatible with most common labeling approaches. Together with the community we will extend the collection of Nup96 cell lines to other fluorescent proteins and peptide tags. We expect that they will find widespread use in many labs for optimization, quality control and counting and that they become the gold standard to quantify effective labeling efficiencies of new dyes and labels.

## Acknowledgements

PA-JF549, PA-JF646 and Halo-Cy5 were a kind gift of Luke Lavis, HHMI Janelia Research Campus. This work was supported by the European Research Council (ERC CoG-724489, J.R., M.M., P.H., J.V.T.), the National Institutes of Health Common Fund 4D Nucleome Program (Grant U01 EB021223 / U01 DA047728 to J.E. and J.R.), the UK Biotechnology and Biological Sciences Research Council (BB/M022374/1; BB/P027431/1; BB/R000697/1; BB/S507532/1, R.H. and P.M.P.), the Wellcome Trust (203276/Z/16/Z, R.H. and P.M.P.), the EMBL Interdisciplinary Postdoc Programme (EIPOD) under Marie Curie Actions COFUND (Y.L.) and the European Molecular Biology Laboratory (J.V.T., K.C., P.H., S.K.P., K.K., Y.L., Y.W., M.M., U.M., B.N., M.K., V.J.S., J.E. and J.R.). V.J.S. acknowledges support by the Boehringer Ingelheim Fonds. Requests for cell lines should be addressed to Jan Ellenberg.

## Author contributions

J.R. conceived the approach, B.N., M.K., V.J.S., J.E., J.V.T. and U.M. generated the cell lines, J.V.T., M.K., K.C., P.H., S.K.P., M.R., D.H., K.C.K., S.J.H., Y.L., Y.W., M.M., U.M. and J.R. developed the methods, wrote the software, acquired and analyzed the data. R.H. and P.M.P. acquired the expansion microscopy data, J.V.T., M.K., P.H., M.M. and J.R. wrote the manuscript with input from all authors.

## Competing financial interests

The authors declare no competing financial interests.

## Methods

### Generation of CRISPR cell lines

Genome editing was performed using CRISPR-Cas9D10A nickase as described in Koch et al.^19^ The gRNA sequences for Nup96 C-term are as follows, sense: 5’-GTTGGGAGCCTGTGAGCCCC-3’ and antisense: 5’-CAGTTCTCGCAGATAGGACT-3’.

The synthetic gene pNup96-mEGFPDonor plasmid encoding for left (1.1 kb) and right (0.8 kb) homology arms for the C-terminus of Nup96 was assembled from synthetic oligonucleotides and/or PCR products. A linker sequence (5’ ACTAGTCGACGGTACCGCGGGCCCGGGATCCACCGGCCGGTCGCCACC 3’) between the left homology arm containing multiple cloning sites was inserted to aid the generation of donor plasmids encoding for other tags. The fragment was inserted into the pMA-RQ (ampR) vector backbone.

Donor plasmids encoding for mMaple^20^, SNAP_f_ tag^54^ (NEB) and HaloTag (Promega) were generated by swapping out mEGFP using restriction enzymes *EagI-HF* and *NheI-HF* (NEB).

#### Southern blotting of Nup96

Southern blotting was performed in accordance Koch et al.^19^ Genomic DNA was prepared using the Wizard Genomic DNA Purification kit (Promega) and digested with *SspI-HF* and *MfeI-HF* (NEB). The probe sequences used are as follows:

Nup96 C-term: (5’-TCCAGTTTCTCTCTGCCACATCCACCTGTTTAAATTATCTACATGGCTTGTGATTTTTCAGGATTTATTACTGTTTTGTGTTTTCTTATTTATTTTCTATCAGTTTCATGAGAGCAAATAACCTGTCTTGCTCTTGATCCTCCTGCCCCCTGCACACAGCTTTTTTGGTGTTTTAGAAAAGGCTATAAACTTGGAGTCAGGGGACCT-3’);
mEGFP: (5’-CACATGAAGCAGCACGACTTCTTCAAGTCCGCCATGCCCGAAGGCTACGTCCAGGAGCGCACCATCTTCTTCAAGGACGACGGCAACTACAAGACCCGCGCCGAGGTGAAGTTCGAGGGCGACACCCTGGTGAACCGCATCGAGCTGAAGGGCATCGACTTCAAGGAGGACGGCAACATCCTGGGGCACAAGCTGGAGTACAACTACAACAGCCACAACGTCTATATCATGGCCGACAAGCAGAAGAACGGCATCAAGGTGAACTTCAAGATCCGCCACAACATCGAGGACGGCAGCGTGCAGCTCGCCGACCACTACCAGCAGAACACCC-3’);
mMaple: (5’-AGCATGACCTACGAGGACGGCGGCATCTGCATCGCCACCAACGACATCACAATGGAGGAGGACAGCTTCATCAACAAGATCCACTTCAAGGGCACGAACTT-3’);
SNAPtag: (5’-AAAGACTGCGAAATGAAGCGCACCACCCTGGATAGCCCTCTGGGCAAGCTGGAACTGTCTGGGTGCGAACAGGGCCTGCACCGTATCATCTTCCTGGGCAAAGGAACATCT-3’);
HaloTag: (5’-TGCATTGCTCCAGACCTGATCGGTATGGGCAAATCCGACAAACCAGACCTGGGTTATTTCTTCGACGACCACGTCCGCTTCATGGATGCCTTCATCGAAGC-3’)

#### siRNA silencing of Nup96 in U2OS

To test specificity of the anti-Nup98 antibody, U2OS cells were seeded onto a 35 mm cell culture dish. 48 h after seeding, MISSION^®^ esiRNA Human nup98 (esirna1) (Sigma, EHU087381-20ug, Lot: BEV) was introduced using lipofectamine 2000 (life technologies). 48 h after transfection the cell layer was scrapped and cell lysate was collected for western blot analysis.

#### Western blotting of Nup96

U2OS cell lysates were collected in Pierce RIPA buffer (Cat#89900; Lot no. NF170965; ThermoFisher Scientific) supplemented with Complete protease inhibitors (Roche) and phenylmethanesulfonylfluoride (PMSF). Cell lysate protein concentration was determined using Pierce BCA protein assay kit (Cat#23225; Lot no. QI223168; ThermoFisher Scientific). 50 μg of cell lysate was loaded onto a 4-12% gradient gel and ran at 165 V constant for 45-60 min in 1X MOPS-SDS buffer (NuPAGE) at room temperature (RT). Proteins were then transferred to a PVDF membrane at 15 V constant for 60 min in cold 1X transfer buffer supplemented with 10% (v/v) methanol (Bolt™) at RT. Membranes were then blocked in 10% (w/v) milk in TBS-T pH 7.6 for 1 hour at RT. After blocking, membranes were incubated in 1:2000 diluted primary antibody (pAb anti-Nup98, Cat#NB1000-93325; LotA1; Novus) in 3% (w/v) BSA in TBS-T at 4 °C overnight. Membranes were then incubated in 1:10000 diluted secondary antibody in 5% (w/v) milk in TBS-T for 1 hour at RT. Chemiluminescence reagents were added to the membrane with subsequent film exposure.

### Sample preparation

#### Buffers

**Table 2:** Buffers used in this work

| Buffer | Composition | Reference |
| --- | --- | --- |
| **FB**<br>Fixation buffer | 2.4% (w/v) formaldehyde in PBS |  |
| **PB**<br>Permeabilization buffer | 0.4% (v/v) Triton X-100 in PBS |  |
| **QS**<br>Quenching solution | 100 mM NH<sub>4</sub>Cl in PBS |  |
| **TRB**<br>Transport buffer | 20 mM HEPES pH 7.5<br>110 mM KAc<br>1 mM EGTA<br>250 mM Sucrose<br>in H<sub>2</sub>O | Pleiner et al., 2015<sup>55</sup><br>Göttfert et al., 2017<sup>56</sup> |
| **TBA**<br>Transport buffer with BSA | 1% (w/v) BSA<br>in **TRB** | Pleiner et al., 2015<sup>55</sup><br>Göttfert et al., 2017<sup>56</sup> |

#### Sample seeding

Prior to seeding of cells, high-precision 24 mm round glass coverslips (No. 1.5H; Cat#117640; Marienfeld, Lauda-Königshofen, Germany) were cleaned by placing them overnight in a methanol/hydrochlorid acid (50/50) mixture while stirring. Following that, the coverslips were repeatedly rinsed with water until they reached a neutral pH. They were then placed overnight into a laminar flow cell culture hood to dry them before finalizing the cleaning of coverslips by UV-irradiation for 30 min.

For superresolution microscopy, homozygous endogenously tagged cells were seeded on clean glass coverslips two days prior fixation in such a way, that they reach a confluency of about 50-70% on the day of fixation. For diffraction limited techniques, cells were seeded on 35 mm cell culture dishes with a 10 mm glass bottom insert (Cat#627860; Greiner Bio-One) instead. Cells grew on the coverslip or the 35 mm cell culture dish in growth medium (DMEM [Gibco; #11880-02] containing 1x MEM NEAA [Cat#11140-035; Gibco], 1x GlutaMAX [Cat#35050-038; Gibco] and 10% [v/v] fetal bovine serum [Cat#10270-106; Gibco]) for approximately two days at 37 °C and 5% CO_2_. Before further processing, the growth medium was aspirated, and samples were rinsed two times with PBS to remove dead cells and debris.

#### Expansion microscopy (proExM)

Expansion of samples was performed as described elsewhere^57^. Briefly, monomer solution (1x PBS, 2 M NaCl, 8.625% [w/w, Sigma] sodium acrylate, 2.5% [w/w, Sigma] acrylamide, 0.15% [w/w, Sigma] *N,N*′-methylenebisacrylamide) was mixed and cooled to 4 °C before use. Ammonium persulfate (APS, BIORAD) initiator and tetramethylethylenediamine (TEMED, Sigma) accelerator were added to the monomer solution up to 0.2% (w/w) each. Samples on coverslips were incubated with the monomer solution plus APS/TEMED in a humidified 37 °C incubator for 1 h for gelation. Proteinase K (New England Biolabs) was diluted 1:100 to 8 units/mL in digestion buffer (50 mM Tris/HCl pH 8, 1 mM EDTA, 0.5% [v/v] Triton X-100, 1 M NaCl, Sigma) and incubated with the gels fully immersed in proteinase solution overnight at 23 °C. Digested gels were next placed in excess volumes of double deionized water for 3−4 h to expand (water changed every 30 min), until the size of the expanding sample plateaued. A small piece of the expanded sample was mounted in an ATTOFLUOR chamber (ThermoFisher Scientific) on 18 mm PLL (Sigma) coated coverslips (Marienfeld) and covered with low-melting agarose (Sigma). To determine the level of sample expansion, the average size of nuclei pre- and post-expansion was measured.

#### Nanobody labeling of Nup96-mEGFP fusion proteins

U2OS-Nup96-mEGFP cells, either prepared on glass coverslips for superresolution measurements or 35 mm cell culture dishes for diffraction limited techniques, were stained according to a protocol previously described by Pleiner and colleagues^55^. For this, samples were prefixed for 30 s in **TRB** containing 2.4% (w/v) formaldehyde (FA), followed by washing twice in **TRB** for 5 min each. Plasma membrane-specific permeabilization was achieved by 8 min incubation on ice in **TRB** containing 25 μg/mL digitonin (Cat#D141; Sigma Aldrich). Samples were washed twice for 5 min in **TBA**. First round of staining was achieved by incubating the samples upside-down in a drop of **TBA** containing 100 nM of anti-GFP nanobodies (NanoTag Biotechnologies, FluoTag-Q [Cat#N0301] or FluoTag-X4 [Cat#N0304], either conjugated to AF647, CF680 or STAR 635P) for 30 min on ice. Residual nanobodies were rinsed away in **TBA** twice for 5 min each before cells were further fixed in **TBA** containing 3% (w/v) FA for 10 min followed by two additional washing steps in **TBA** for 5 min each. Permeabilization of the nuclear envelope was facilitated by 3 min incubation in **PB**. Samples were washed twice in PBS for 5 min each before exposing them again upside-down onto a drop of anti-GFP nanobodies (50 nM in **TBA**, same nanobodies as in the first round of staining) for 30 min on ice. Finally, weakly bound and unbound nanobodies were rinsed off in PBS twice for 15 min. For STED-imaging, FluoTag-X4-STAR 635P stained samples were mounted upside-down on glass microscopy slides (ThermoFisher Scientific) using Mowiol (Calbiochem). Edges were further sealed by nail polish and then dried overnight at RT.

#### HaloTag labeling of fixed cells

U2OS-Nup96-Halo cells were stained on previously prepared coverslips using a slightly modified version of the nanobody labeling protocol described above^55^. Instead of staining the samples in two separate rounds of nanobodies (100 nM in round 1 and 50 nM in round 2), the samples were incubated in HaloTag dye buffer (5 μM of Cy5-HaloTag ligand [Lavis Lab, HHMI Janelia Research campus] or HaloTagligand-O2-AF647/HaloTag-ligand-O4-AF647 [custom substrates from Peps4LS, Heidelberg] in **TBA**) for 1 h at RT in both incubation steps. All other steps were performed in accordance to the above described protocol.

#### HaloTag live labeling

Coverslips covered in an approximately 50-70% confluent layer of U2OS-Nup96-Halo were incubated in pre-warmed growth medium containing PA-JF549 or PA-JF646-HaloTag ligand (250 – 5000 nM were tested without significant difference in labeling efficiency; Lavis Lab, HHMI Janelia Research campus) at 37 °C and 5% CO_2_ for 1 h. The samples were subsequently rinsed thrice in pre-warmed PBS and incubated in pre-warmed growth medium without dye for 1 h at 37 °C and 5% CO_2_ to wash off non-covalently bound dye. Following that, the samples were rinsed three times in PBS before prefixing them at RT for 30 s in **FB**. Permeabilization was facilitated in **PB** for 3 min before completing the fixation process for 30 min in **FB**. Subsequently, FA was quenched by incubating the coverslip for 5 min in **QS**. Sample preparation was finalized by washing twice in PBS for 5 min each.

#### SNAP-tag labeling of fixed cells

U2OS-Nup96-SNAP cells were prefixed for 30 s in **FB** before permeabilization in **PB** for 3 min. To complete fixation, samples were incubated for 30 min in **FB**. FA was subsequently quenched in **QS** for 5 min before washing the coverslip twice for 5 min in PBS. To reduce unspecific binding, the sample was incubated for 30 min with Image-iT FX Signal Enhancer (ThermoFisher Scientific) before staining in SNAP dye buffer (1 μM BG-AF647 [New England Biolabs; #S9136S], 1 μM DTT in 0.5% [w/v] BSA in PBS) for 2 h at RT. To remove unbound dye, coverslips were washed three times in PBS for 5 min each.

#### Fixation of mMaple tagged cell lines

Glass coverslips prepared with U2OS-Nup96-mMaple or HEK-Nup107-mMaple cells were prefixed for 30 s in **FB** before incubation in **PB** for 3 min. To complete fixation, samples were incubated for 30 min in **FB**. FA was subsequently quenched in QS for 5 min before washing the coverslip twice for 5 min in PBS.

#### Strain & sample preparation for yeast

For protein counting in *Saccharomyces cerevisiae*, the respective proteins (Nup188, Nup82, Nup192, Nic96, Nup49 and Abp1) were endogenously tagged on the C-terminus by homologous recombination. Shortly, we constructed plasmids encoding mMaple and different selectable markers by standard molecular biology methods^58^. The cassette containing a peptide linker, mMaple and the selectable marker was amplified by PCR and transformed into competent yeast cells. Yeast cells were plated on selective plates, grown for 2-3 days until single colonies were obtained. Correct tagging was confirmed by colony PCR and imaging.

N-terminal labeling of Nic96 was performed seamlessly^59^. First, a cassette was amplified by PCR from a vector that contains the first 180 bp of mMaple, the selectable marker for the expression of the URA3 gene and a promoter for the tagged gene of interest surrounded by two I-*Sce*I restriction sites and full-length mMaple. This cassette was transformed into yeast cells that express I-*SceI* under control of a galactose inducible promotor. After correct integration was confirmed by colony PCR, the strain was cultivated on plates containing galactose to induce the expression of I-*Sce*I and resistance cassette loopout. Successful excision was counterselected on plates containing 5-fluoroorotic acid.

For immobilization of yeast, the coverslips were coated with concanavalin A (ConA; Cat#C2010; Sigma-Aldrich). For this, the coverslips were cleaned overnight in a 1:1 mixture of methanol and hydrochloric acid, washed 3 times with dH_2_O and plasma-cleaned. Next, 20 μl of 4 mg/mL ConA in PBS was pipetted onto the coverslip and spread, incubated under a humidified atmosphere for 30 min and then dried.

For super-resolution imaging, the respective strains were grown at 30 °C shaking at 220 rpm in synthetic complete medium without tryptophan (SC-Trp) to reduce autofluorescence. A 4 mL overnight culture was inoculated from a single colony on a freshly restreaked plate. In the morning of the experiment, the culture was diluted to an optical density (OD_600_) of 0.25 in 10 mL of SC-Trp medium and cultured for approximately 3 more hours to logarithmic phase. Then, cells from the reference strain (Nup188-mMaple Abp1-mEGFP) and the respective target strain were mixed in a 1:1 ratio and spinned down in a table top centrifuge (3 min at 1000 g and RT), resuspended in about 200 μl of residual medium and pipetted onto a ConA-coated coverslip. All subsequent steps were carried out in the dark to prevent pre-conversion of mMaple. After allowing the cells to settle for 20 min, the coverslips were fixed in fixation solution (4% [w/v] FA, 2% [w/v] sucrose, in PBS) for 15 min. Subsequently, remaining FA was quenched by washing twice for 5 min in **QS**. After washing 3 time with PBS for 5 min each, the coverslip was ready for imaging. The sample was mounted on a custom sample holder in imaging buffer (50 mM Tris pH 8 in 95% [v/v] D_2_O) and subjected to SMLM.

**Table 3:** List of yeast strains used in this study

| Strain | Genotype | Source |
| --- | --- | --- |
| MKY0100 | MATa, his3Δ200, leu2-3,112, ura3-52, lys2-801 | Kaksonen lab |
| MKY0122 | MATa, his3Δ200, leu2-3,112, ura3-52, lys2-801, ABP1-mEGFP::HIS3MX6 | Kaksonen lab |
| Nup188-mMaple | MATa, his3Δ200, leu2-3,112, ura3-52, lys2-801, NUP188-mMaple::HIS3MX6 | This study |
| Nup82-mMaple | MATa, his3Δ200, leu2-3,112, ura3-52, lys2-801, NUP82-mMaple::HIS3MX6 | This study |
| Nup192-mMaple | MATa, his3Δ200, leu2-3,112, ura3-52, lys2-801, NUP192-mMaple::HIS3MX6 | This study |
| mMaple-Nic96 | MATa, his3Δ200, leu2-3,112::GalL-ISce-natNT2, ura3-52, lys2-801, mMaple-NIC96 | This study |
| Nic96-mMaple | MATa, his3Δ200, leu2-3,112, ura3-52, lys2-801, NIC96-mMaple::HIS3MX6 | This study |
| Abp1-mEGFP<br>Nup188-mMaple | MATa, his3Δ200, leu2-3,112, ura3-52, lys2-801, ABP1-mEGFP::HIS3MX6, NUP188-mMaple::hphNT1 | This study |
| mMaple-Nic96<br>Nup49-mEGFP | MATa, his3Δ200, leu2-3,112::GalL-ISce-natNT2, ura3-52, lys2-801, mMaple-NIC96, NUP49-mEGFP::HIS3MX6 | This study |
| Nup188-mMaple<br>Nup82-mMaple | MATa, his3Δ200, leu2-3,112, ura3-52, lys2-801, NUP188-mMaple::HIS3MX6, NUP82-mMaple::LEU2 | This study |

### Microscopy

#### Microscope setup and imaging

All SMLM data were acquired on a custom built widefield setup described previously^5,60^. Briefly, the free output of a commercial laser box (LightHub; Omicron-Laserage Laserprodukte, Dudenhofen, Germany) equipped with Luxx 405, 488 and 638 and Cobolt 561 lasers and an additional 640 nm booster laser (iBeam smart, Toptica, Gräfelfing, Germany) were collimated and focused onto a speckle reducer (Cat#LSR-3005-17S-VIS; Optotune, Dietikon, Switzerland) before being coupled into a multi-mode fiber (Cat#M105L02S-A; Thorlabs, Newton, NJ, USA). The output of the fiber was magnified by an achromatic lens and focused into a sample to homogeneously illuminate an area of about 1000 μm^2^. Alternatively, a single-mode fiber (Omicron, LightHUB) could be plugged into the output of the laserbox to allow TIRF imaging. The laser is guided through a laser cleanup filter (390/482/563/640 HC Quad; AHF, Tübingen, Germany) to remove fluorescence generated by the fiber. Emitted fluorescence was collected through a high-numerical-aperture (NA) oil-immersion objective (160x/1.43-NA; Leica, Wetzlar, Germany), filtered by a bandpass filter (525/50 [Cat#FF03-525/50-25, Semrock, Rochester, NY, USA] for mEGFP; 600/60 [Cat#NC458462, Chroma, Bellows Falls, VT, USA] for mMaple and PA-JF549 and 700/100 [Cat#ET700/100m, Chroma] for AF647, Cy5, PA-JF646 and CF680) and imaged onto an Evolve512D EMCCD camera (Photometrics, Tucson, AZ, USA). The z focus was stabilized by an IR-laser that was totally internally reflected off the coverslip onto a quadrant photodiode, which was coupled into closed-loop feedback with the piezo objective positioner (Physik Instrumente, Karlsruhe, Germany). Laser control, focus stabilization and movement of filters was performed using a field-programmable gate array (Mojo; Embedded Micro, Denver, CO, USA). The pulse length of the 405 nm (laser intensity 27.5 W/cm^2^) laser is controlled by a feedback algorithm to sustain a predefined number of localizations per frame. Typical acquisition parameters can be found in **Table 4**. Coverslips containing prepared samples were placed into a custom build sample holder and 500 μL of suitable buffer, depending on the used cell line and experiment (**Table 5**), was added. To avoid a pH drift caused by accumulation of glucuronic acid in GLOX-buffers, the buffer solution was exchanged after about 2 h of imaging. Samples were imaged until close to all fluorophores were bleached and no further localizations were detected under continuous UV irradiation.

**Table 4:**
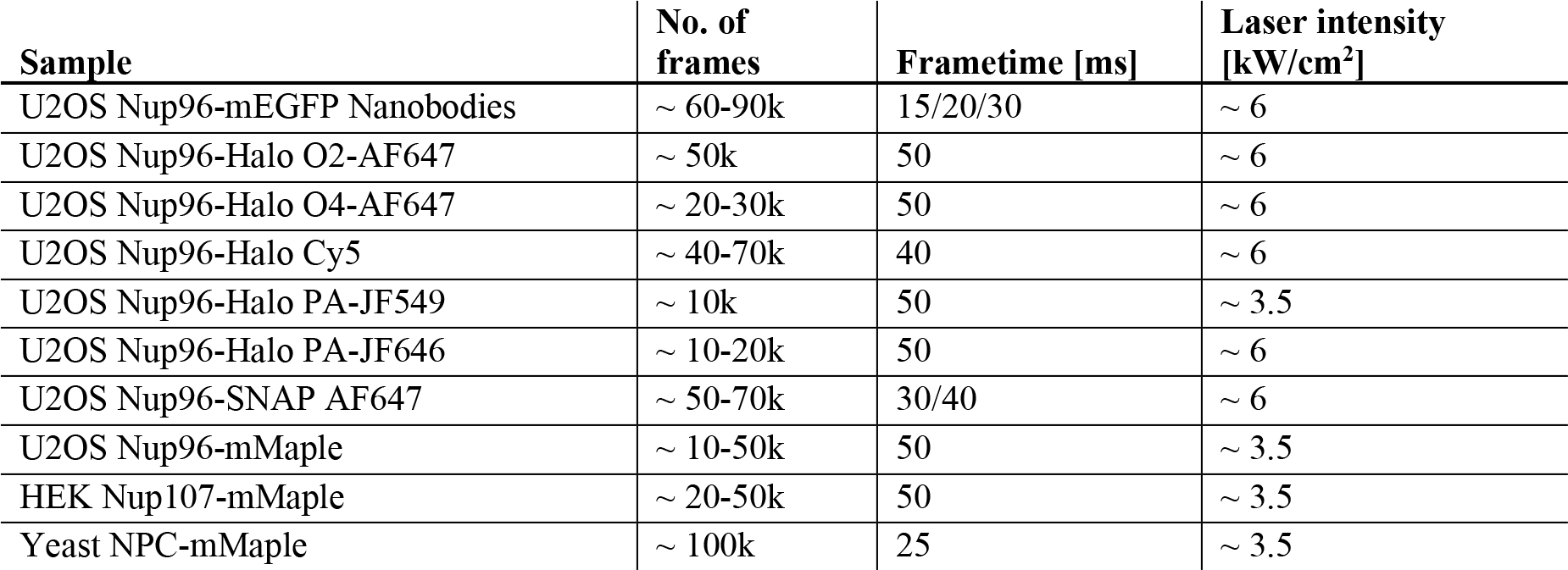
Acquisition parameters during SMLM imaging.

**Table 5:** Used imaging buffers.

| Buffer | Composition | Samples | Reference |
| --- | --- | --- | --- |
| 50 mM Tris in D<sub>2</sub>O | 50 mM Tris/HCl pH 8<br>in 95% (v/v) D<sub>2</sub>O | U2OS Nup96-Halo PA-JF549<br>U2OS Nup96-Halo PA-JF646<br>U2OS Nup96-mMaple<br>HEK Nup107-mMaple<br>Yeast NPC-mMaple | Ong et al., 2015<sup>61</sup> |
| GLOX/MEA | 50 mM Tris/HCl pH 8<br>10 mM NaCl<br>10% (w/v) D-Glucose<br>500 µg/mL Glucose oxidase<br>40 µg/mL Glucose catalase<br>35 mM MEA<br>in H<sub>2</sub>O | U2OS Nup96-mEGFP Nanobodies<br>U2OS Nup96-Halo O2-AF647<br>U2OS Nup96-Halo O4-AF647<br>U2OS Nup96-Halo Cy5<br>U2OS Nup96-SNAP AF647 | Heilemann et al., 2005<sup>62</sup> |
| GLOX/BME | 50 mM Tris/HCl pH 8<br>10 mM NaCl<br>10% (w/v) D-Glucose<br>500 µg/mL Glucose oxidase<br>40 µg/mL Glucose catalase<br>143 mM BME<br>in H<sub>2</sub>O | U2OS Nup96-SNAP AF647 | Bates et al., 2005<sup>63</sup> |
| GLOX/MEA in D<sub>2</sub>O | 50 mM Tris/HCl pH 8<br>10 mM NaCl<br>10% (w/v) D-Glucose<br>500 µg/mL Glucose oxidase<br>40 µg/mL Glucose catalase<br>35 mM MEA<br>in 90% (v/v) D<sub>2</sub>O | U2OS Nup96-SNAP AF647 | Klehs et al., 2014<sup>37</sup> |
| Sulfite | 50 mM Tris/HCl pH 8<br>50 mM Na<sub>2</sub>SO<sub>3</sub>/NaOH pH 8<br>35 mM MEA<br>in H<sub>2</sub>O | U2OS Nup96-SNAP AF647 | Hartwich et al., 2018<sup>39</sup> |

#### Pixel size calibration

The effective pixel size of the microscope was calibrated by translating fluorescent beads, immobilized on a coverslip, with a calibrated sample stage (SmarAct, Oldenburg, Germany) that operated in close loop. From the measured translation of many beads the pixel size could be calibrated with a high accuracy.

#### Widefield, SIM and SRRF on Expanded Samples

After expansion (protocol described above) U2OS-Nup96-mEGFP cells labeled with an Atto488-coupled anti-GFP nanobody were imaged in a Zeiss Elyra PS.1 system. An 100x TIRF objective (Plan-APOCHROMAT 100 /1.46 Oil, Zeiss) was used, with additional 1.6 magnification, to collect fluorescence onto an EMCCD camera (iXon Ultra 897, Andor), yielding a pixel size of 100 nm. Sample was illuminated with a 488 nm laser set at 150 mW/cm^2^. Widefield images were collected with 100 ms exposure, SIM images with 100 ms exposure and 5 grid roations, each SRRF image was generated from a frame-burst of 100 images acquired at 33 Hz. SIM reconstructions were generated with the Zeiss Elyra Zen software using automatic settings. SRRF images were analysed with NanoJ-SRRF^26^ using standard settings. Images were validated for quality using SIMCheck^7^ (SIM) and NanoJ-SQUIRREL^8^ (SIM and SRRF).

#### Confocal microscopy

Fixed U2OS-Nup96-mEGFP samples on 35 mm glass bottom dishes were prepared according to the preparation protocol described above and imaged using an Olympus FV3000 laser scanning microscope. A 60x / 1.40 NA oil immersion objective (Olympus; PLAPON 60XOSC2) was used in combination with a motorized stage, operated by the Fluoview software (Olympus). Pixel size was set to ~ 70 nm in x and y. Fluorescence emission went through a 550/100 bandpass filter and a 1.0 airy unit (202 μm) wide pinhole before detection on 4 GaAsP spectral detectors. For each nucleus, a z-stack, consisting of 3-5 planes 250 nm apart from each other, was acquired around the basal plane of the nucleus to obtain maximum fluorescence intensity for all NPCs.

#### Airy-scan microscopy

35 mm glass bottom dishes containing U2OS-Nup96-mEGFP were fixed in accordance to the previously described protocol. A Zeiss LSM 880 with an additional Airy FAST detector module (Zeiss) was used for airy-image acquisition in combination with a 63x / 1.4 NA oil immersion objective (Zeiss; Plan-Apochromat 63x/1.4 Oil DIC M27). The system was operated by the ZEN software (Zeiss; black edition). Pixel size was set to ~ 40 nm in x and y direction. Samples were focused on the basal plane of the nucleus and mEGFP was excited using a 488 nm laser. Emission was collected through a 495-550 nm bandpass filter, 570 nm longpass filter and a 1.25 airy unit (~60 μm) pinhole onto the 32 GaAsP detector elements. A z-stack, consisting of 3-5 slices 200 nm apart from each other around the basal plane were acquired for each nucleus. Post-processing was done with ZENs airy-scan processing, using automatic deconvolution parameters.

#### STED microscopy

Samples were prepared according to the protocol for nanobody staining of U2OS-Nup96-GFP samples and were imaged on an Abberior STED/RESOLFT microscope (Abberior Instruments; Expert Line) running the Imspector software (Abberior Instruments). The microscope comprises of an IX83 stage (Olympus) in combination with a UPlan-S Apochromat 100x / NA 1.40 oil objective (Olympus). Pixel size was set to 15 nm in x, y direction. Super-resolved images were acquired by donut-shaped depletion using a 775 nm pulsed laser along with a 640 nm pulsed laser, exciting STAR 635P tagged Nup96-mEGFP. A single plane of the lower side of the nucleus was imaged. Emission was collected through a 685/70 nm bandpass filter.

#### Ratiometric dual-color SMLM

For ratiometric dual-color imaging of AF647 and CF680, the emitted fluorescence was split by a 665LP beamsplitter (Cat#ET665lp, Chroma), filtered by a 685/70 (Cat#ET685/70m, Chroma) bandpass filter (transmitted light) or a 676/37 (Cat#FF01-676/37-25, Semrock) bandpass filter (reflected light) and imaged side by side on the EMCCD camera. The color of the individual blinks was assigned by calculating the ratio of the intensities in the two channels.

#### Astigmatic 3D SMLM

3D SMLM data was acquired using a cylindrical lens (f = 1000 mm; Cat#LJ1516L1-A, Thorlabs) to introduce astigmatism. The data were fitted and analyzed as described previously^64^. First, z-stacks with known displacement of several (15-20) fields of view of TetraSpeck beads on a coverslip were acquired to generate a model of the experimental point spread function. This model was then used to determine the z-position of the individual localizations. To correct for depth-dependent aberrations, we acquired stacks of beads in agarose to determine the fitting errors as described previously^31^.

### Data analysis

All data analysis was performed with custom software written in MATLAB and is available as open source (github.com/jries/SMAP). Installation instructions are found in the README.md, and step-by-step guides on how to use the software to perform all analyses used in this manuscript are available via the *Help* menu.

#### Post-processing

x-, y-, and, when applicable, z-positions were corrected for residual drift by a custom algorithm based on redundant cross-correlation. Localizations persistent in consecutive frames were grouped into one localization. Localizations were filtered by the localization precision and the log-likelihood to exclude dim or poorly fitted localizations, and for 2D data by the fitted size of the PSF to exclude localizations that were strongly out-of-focus. Super-resolution images were constructed with every localization rendered as a 2D elliptical Gaussian with a width proportional to the localization precision.

#### Determination of the expansion factor

To determine the local expansion factor from the Ex-SIM data set, we manually selected positions of 203 nuclear pores and fitted a cropped image of the pore with a model that consisted of a ring convoluted with a Gaussian function, treating the radius and the standard deviation of the Gaussian as free fitting parameters. We then re-fitted the data keeping the standard deviation of the Gaussian fixed to its mean value. By comparing the mean value of the radius with that one measured on a calibrated SMLM microscope, we directly determined the expansion factor.

#### Segmentation

To automatically segment nuclear pore complexes, we convoluted the reconstructed superresolution image with a kernel consisting of a ring with a radius corresponding to the radius of the NPC, convoluted with a Gaussian. Local maxima over a user-defined threshold were treated as candidate NPCs. These candidates included many aberrant structures. We cleaned up the segmentation by a two-step filtering process: 1) We fitted the localizations corresponding to each candidate with a circle to reject structures with very small (typically < 40 nm) or very large (>70 nm) radii. 2) We re-fitted the localizations with a circle of fixed radius to determine its center coordinates, and rejected structures where more than 25% of the localizations were within 40 nm of the center (structures that visually did not resemble NPCs) or more than 40% of the localizations were further away than 70 nm (structures that were usually composed of two adjacent NPCs and wrongfully segmented).

We segmented many data sets manually and compared that segmentation with the automatic segmentation and found an excellent agreement with less than 1.2% difference in measured ELE values and less than 5% error in the mean number of localizations per NPC.

#### Geometric analysis

All geometric analysis was performed on NPCs segmented as described above, based on the coordinates of the localizations.

##### Analysis of profiles

Profiles in the x-y plane were constructed by 1) selecting a linear ROI in the direction the profile is calculated, 2) selecting only localizations in a rectangular ROI along the line profile and with a given width, 3) rotating the coordinates such that the x’-axis is along the direction of the line profile, 4) calculating a histogram of the x’ coordinates. This histogram was then fitted with a single or double Gaussian function. For profiles along the z-direction we 1) defined a ROI, 2) calculated the histogram of z-coordinates for localizations within this ROI and 3) fitted the histogram with a single or double Gaussian function.

We want to stress that care must be taken that profiles are constructed from a sufficient number of localizations, and are never measured in a superresolution image where localizations are rendered with a Gaussian kernel. Otherwise even single localizations can result in ‘profiles’ with arbitrary small width and two random localizations can be ‘resolved’ if their distance is larger than the arbitrary kernel size. This holds true for any profile analysis of SMLM data and is not restricted to NPCs.

##### Radius of the NPC

The radius of the circular NPC structures was determined by directly fitting the coordinates of the localizations with a circular model treating the x and y coordinates and the radius as free fitting parameters.

##### Distance between cytoplasmic and nucleoplasmic rings in 2D data

Ring distances were measured on 2D data sets where the focus was set to the mid-plane of the nucleus. 1) We manually segmented structures on vertical parts of the nuclear envelope. 2) We constructed profiles perpendicular to the nuclear envelope with a width of 200 nm by calculating the histogram of rotated localizations. 3) We fitted the profiles with a double-Gaussian function to determine the distance of the rings.

##### Distance of cytoplasmic and nucleoplasmic rings from 3D data

Segmented localizations were fitted in 3D with a template describing two parallel rings with a fixed radius (mean of the radius as measured before) and variable x, y and z positions, rotation angles and distance between the rings. As a validation, we used the fitted rotation angles to rotate the localizations so that all NPCs were aligned and fitted the z-profile with a double Gaussian as described above for 2D data.

##### Azimuthal angle

We determined the azimuthal angle between the cytoplasmic and nucleoplasmic rings from 3D data. 1) we fitted the localizations with a circle to determine its x and y center coordinates. 2) We determined the axial position of the NPC by fitting the z-profile with a double-Gaussian as described above. 3) We separated localizations belonging to the upper and lower ring. 4) We transformed the x, y coordinates to polar coordinates. 5) We constructed histograms of the polar angles. 6) we calculated the auto- and cross-correlation curves of these histograms taking into account the circular boundary conditions. 7) We calculated the average correlation curves for all NPCs. 8) We fitted the average cross-correlation curve with a cosine function of fixed frequency and varying phase. We fitted the offset and amplitude of the trigonometric function by 3^rd^-degree polynomials. We excluded the central 24° from the fit as they contained strong contributions from the re-activation of fluorophores. 9) The azimuthal angle corresponds to the fitted phase of the trigonometric function.

#### Determination of effective labeling efficiencies

To count the number of visible corners in each nuclear pore complex we used the following approach: 1) The segmented and filtered localizations were fitted by a circle of fixed radius corresponding to the mean radius as determined before and coordinates were converted into polar coordinates *ϕ*_*i*_, *r*_*i*_. 2) Localizations too close to the center of the ring (*r*_*i*_ < 30 nm) or too far away (*r*_*i*_ > 70 nm) were excluded as background localizations. 3) We determined the rotation of the structure by minimizing

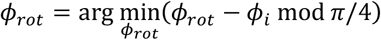

4) we counted the number of segments containing a localization from a histogram of *ϕ*_*i*_ with a bin width of *π*/4 and a start bin of *ϕ*_*rot*_ − *π*/8. 5) We constructed a histogram of the number of corners of all NPCs in the data set and fitted it using the probabilistic model as described below, using the effective labeling efficiency as a free fitting parameter. 6) To calculate the statistical error, we used bootstrapping with typically 20 re-sampled data sets.

##### Probabilistic model for effective labeling efficiency

The binomial probability density function

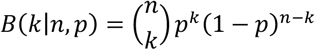

describes the probability of observing *k* successes in *n* independent trials, where the probability of success in any given trial is *p*. Thus, the probability of a corner of the NPC (consisting of 4 labels) to be dark is *p*_dark_ = *B*(0|4, *p*_label_) and the probability to see a corner with at least one label is *p*_bright_ = 1 − *p*_dark_. The probability of *N* out of 8 corners being bright and visible is:

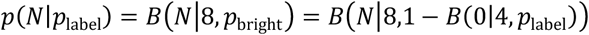

##### Determination of number of localizations per fluorophore

The number of localizations (blinking events) *N*_*b*_ that are detected per fluorophore can be directly calculated from the ELE, the number of localizations per NPC *N*_*l*_ and the number of Nup96 molecules per NPC *N*_Nup96_ = 32:

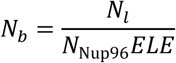

#### Simulations

To validate our analysis routines, we performed realistic simulations based on a two-state (bright and dark) fluorophore model with bleaching^9^: 1) We defined the 3D coordinates of the 32 Nup96 proteins in the nuclear pore complex based on our calibration (Figure 2). 2) we randomly displace all coordinates by a random vector and rotate the coordinates in 3D by random angles. 2) With a probability *p*_label_ a protein is labeled and creates a localization. 3) A labeled protein has a probability *p*_react_ to be reactivated. 4) Whenever a fluorophore is activated it appears at random during a frame and lives for *t*_*l*_ frames, determined as a random variable from an exponential distribution. 5) When it is on, a fluorophore has a constant brightness. 7) The emitted photons in each frame are determined as a random Poisson variable with a mean corresponding to the average brightness during the frame. 8) For each frame we calculate the CRLB in x, y and z from the number of photons and the background^65^. 9) This error is added to the true x, y and z positions of the fluorophores as normally distributed random values with a variance corresponding to the respective calculated CRLB.

The simulated localizations were processed with the same data analysis pipeline as the real data.

#### Counting of protein copy numbers

##### Counting in diffraction-limited microscopy using Nup96-mEGFP as a reference

We used a simple data analysis procedure to compare the brightness of reference and target structures in confocal images: 1) We subtracted the image offset, and if required corrected the images for photobleaching. 2) We calculated the maximum intensity projection of 3 frames around the focal plane of the nuclear pore structures and convolved the image with a Gaussian (*σ* = 0.5 pixels). 3) We up-sampled the image by a factor of two using cubic spline interpolation. 4) We determined all local maxima and chose a threshold based on the histogram of intensity values of those maxima. 5) We fitted the histogram of maxima intensities above the threshold with a Gaussian function to determine a robust estimate of the mean of the intensity values 〈*I*_*t*_〉 and 〈*I*_*r*_〉 for reference and target cell lines. 6) With *N*_*r*_ copies of the reference protein in the complex, the copy number in the target complex is then *N*_*t*_ = *N*_*r*_ 〈*I*_*t*_〉/〈*I*_*r*_〉.

##### Counting in mammalian cells using Nup96-mMaple as a reference

1) We automatically segmented reference and target data as described above and only considered nuclear pores in the focus (mean value of the fitted size of the PSF smaller than 145 nm). 2) We counted the number of grouped localizations (*L*_*r*_, *L*_*t*_) in a circular ROI of a diameter of 220 nm. 3) From the mean number of localizations per nuclear pore complex 〈*L*_*t*_〉 and 〈*L*_*r*_〉 we can calculate the copy number of the target complex *N*_*t*_ = *N*_*r*_ 〈*L*_*t*_〉/〈*L*_*r*_〉.

##### Counting in yeast cells using Nup188 as a reference

1) We manually segmented NPCs in yeast cells and excluded structures that were out-of-focus, at the edge of the nucleus or too close to other structures. 2) Based on the intensity of Abp1-mEGFP in a diffraction limited channel we assigned all NPCs in a cell to belong to the reference cell line (significant mEGFP signal) or to the target cell line (no mEGFP signal). 3) We determined the number of localizations in a circular ROI of a diameter of 150 nm. 4) As above, we determined the mean number of grouped localizations and from those the copy number of the target complex.

##### Model to estimate steady state maturation fraction

Here we derive a very simple model to estimate the fraction of matured photoconvertible fluorescent protein (e.g. mMaple) in the steady state neglecting degradation. *P* denotes the amount of not yet matured protein, *M* the amount of the matured protein and *k*_*m*_ is the maturation rate. We assume exponential growth (growth rate *k*_*g*_) of the organism and thus of the proteins:

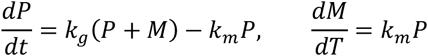

The solution is:

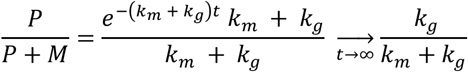

Assuming a doubling time for yeast of 120 min and a maturation time for mMaple of 48 min we find that in the steady state on average 28% of the mMaple is not yet matured. For mammalian cells (generation time 1 day) the fraction is reduced to 3.2%.

## Supplementary Table and Figures

**Supplementary Table 1:** Imaging conditions and analysis.

| Sample | Buffers | ELE [%] | Photons per localization | Localizations per NPC | Localizations per fluorophore | ROIs imaged / NPCs analyzed |
| --- | --- | --- | --- | --- | --- | --- |
| U2OS-Nup96-SNAP-BG-AF647 | GLOX/MEA | 60 ± 3 | 10674 ± 833 | 85 ± 18 | 4.4 ± 0.7 | 8 / 3318 |
| U2OS-Nup96-SNAP-BG-AF647 | GLOX/MEA in D<sub>2</sub>O | 56 ± 3 | 9986 ± 365 | 65 ± 8 | 3.6 ± 0.4 | 5 / 3379 |
| U2OS-Nup96-SNAP-BG-AF647 | GLOX/BME | 63 ± 4 | 12815 ± 824 | 112 ± 10 | 5.6 ± 0.5 | 8 / 3724 |
| U2OS-Nup96-SNAP-BG-AF647 | Sulfite | 39 ± 2 | 6984 ± 611 | 19 ± 2 | 1.5 ± 0.1 | 5 / 2708 |
| U2OS-Nup96-mMaple fixed | 50 mM Tris in D<sub>2</sub>O | 58 ± 4 | 2159 ± 259 | 50 ± 6 | 2.7 ± 0.2 | 23 / 10180 |
| U2OS-Nup96-mMaple live | 50 mM Tris in D<sub>2</sub>O | 57 ± 3 | 1877 ± 50 | 51 ± 1 | 2.8 ± 0.1 | 4 / 1081 |
| U2OS-Nup96-Halo-Cy5 | GLOX/MEA | 38 ± 5 | 11014 ± 722 | 56 ± 13 | 4.6 ± 0.7 | 14 / 5967 |
| U2OS-Nup96-Halo-O2-AF647 | GLOX/MEA | 36 ± 3 | 15530 ± 593 | 39 ± 8 | 3.3 ± 0.4 | 5 / 1393 |
| U2OS-Nup96-Halo-O4-AF647 | GLOX/MEA | 30 ± 1 | 14716 ± 906 | 41 ± 4 | 4 ± 0.3 | 3 / 793 |
| U2OS-Nup96-Halo-PA-JF549 | 50 mM Tris in D<sub>2</sub>O | 21 ± 4 | 9295 ± 1094 | 9 ± 2 | 1.3 ± 0.1 | 17 / 4066 |
| U2OS-Nup96-Halo-PA-JF646 | 50 mM Tris in D<sub>2</sub>O | 19 ± 3 | 17116 ± 1296 | 10 ± 2 | 1.7 ± 0.1 | 3 / 762 |
| U2OS-Nup96-mEGFP-NB-Q-AF647 | GLOX/MEA | 59 ± 3 | 6778 ± 570 | 54 ± 8 | 2.8 ± 0.3 | 6 / 2913 |
| U2OS-Nup96-mEGFP-NB-Q-CF680 | GLOX/MEA | 62 ± 1 | 6147 ± 182 | 101 ± 16 | 5.1 ± 0.8 | 5 / 1805 |
| U2OS-Nup96-mEGFP-NB-X4-AF647 | GLOX/MEA | 73 ± 6 | 6207 ± 596 | 91 ± 15 | 3.9 ± 0.4 | 9 / 4303 |
| U2OS-Nup96-mEGFP-NB-X4-CF680 | GLOX/MEA | 75 ± 4 | 6490 ± 256 | 132 ± 20 | 6.4 ± 2.3 | 7 / 2395 |

**Supplementary Figure 1:**
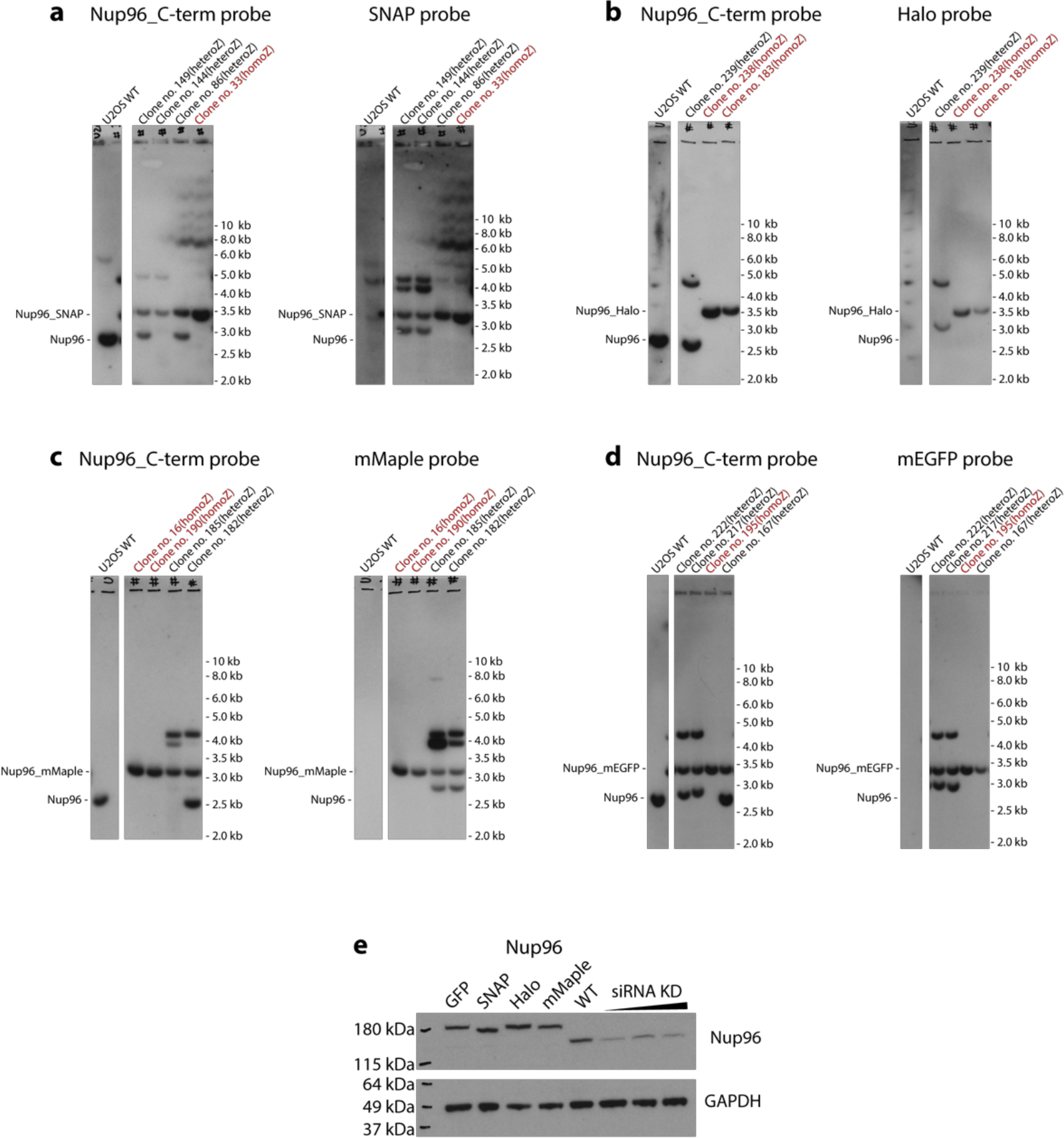
Validation of cell line homozygosity. (**a-d**) Southern blots of (**a**) **Nup96-SNAP,** (**b**) **Nup96-Halo,** (**c**) **Nup96-mMaple,** (**d**) **Nup96-mEGFP.** Blots on the left were generated from probes against Nup96-C-term and blots on the right were generated from probes against respective tags. The presence of a single band was indicative of a homozygous knock-in (in red). (**e**) Western blot of the homozygous cell lines probed with an anti-Nup98 antibody. Reduction of band intensity in Nup96-WT indicates the specificity of the antibody to Nup96. siRNA concentrations used: 0.6 μg, 1.2 μg and 1.8 μg.

**Supplementary Figure 2:**
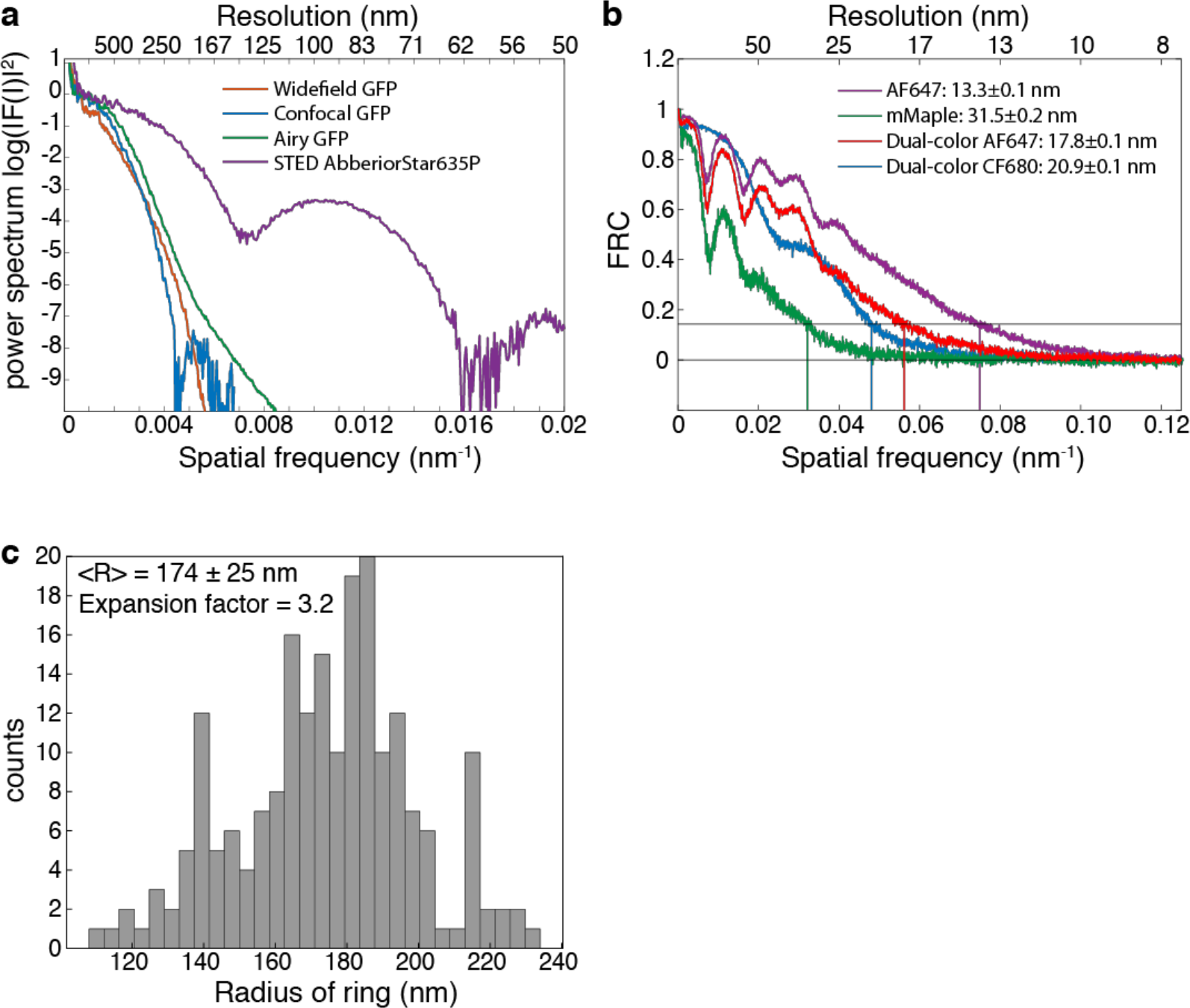
Resolution. (**a**) **Logarithmic power spectrum** for **Fig. 1 f-i**. (**b**) **Fourier Ring Correlation** for **Fig. 1 m-q** including resolution estimates. (**c**) Histogram of radii resulting from a ring fit to **Fig. 1p**. From the average measured radius <R> = 174 ± 25 nm and the known radius (**Fig. 2**) the expansion factor was estimated to be 3.2.

**Supplementary Figure 3:**
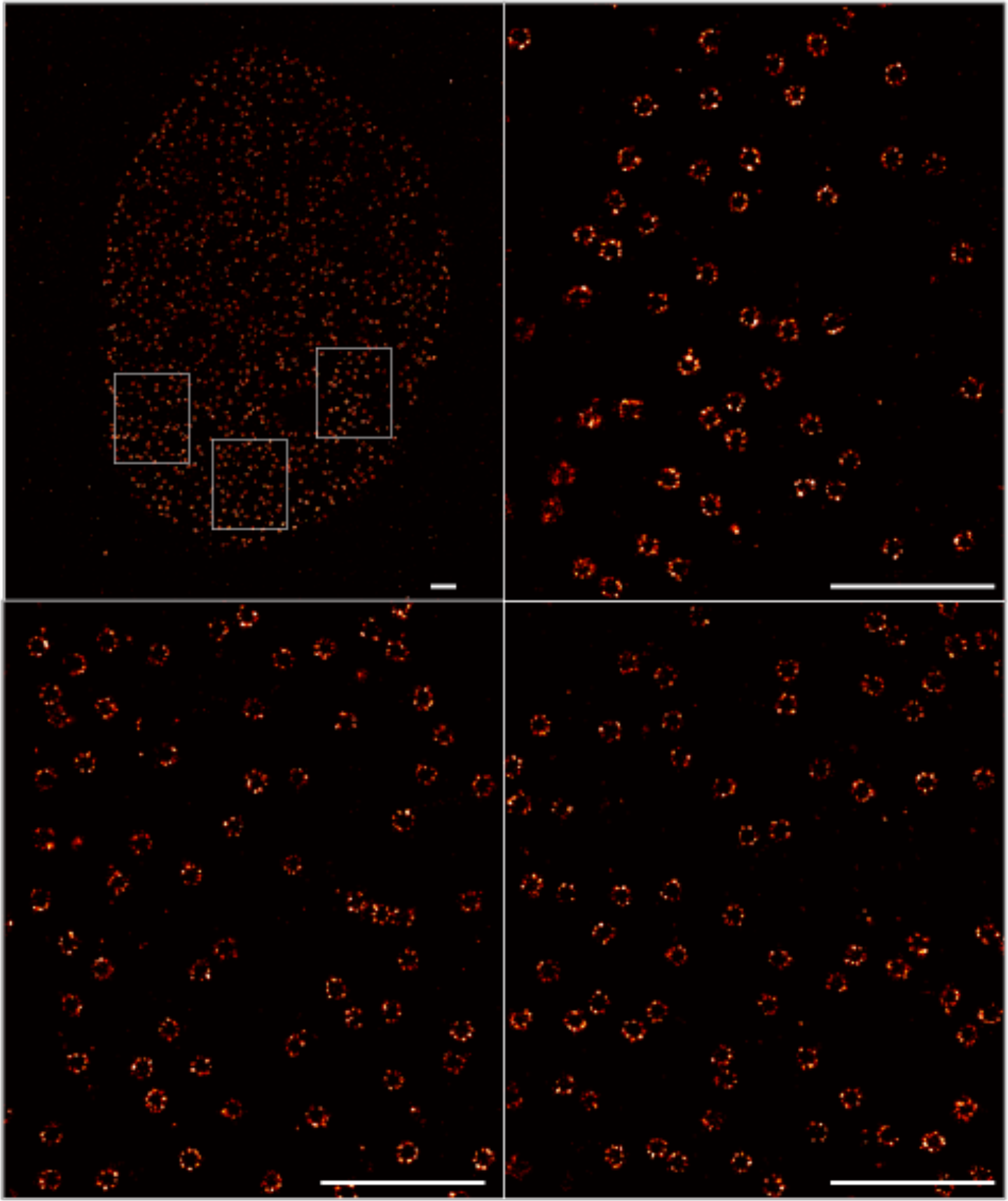
SMLM with total internal reflection fluorescence (TIRF) excitation on Nup96-SNAP-AF647. In the majority of cells the lower nuclear envelope is sufficiently close to the coverslip to be efficiently excited with TIRF. Scale bars 1 μm.

**Supplementary Figure 4:**
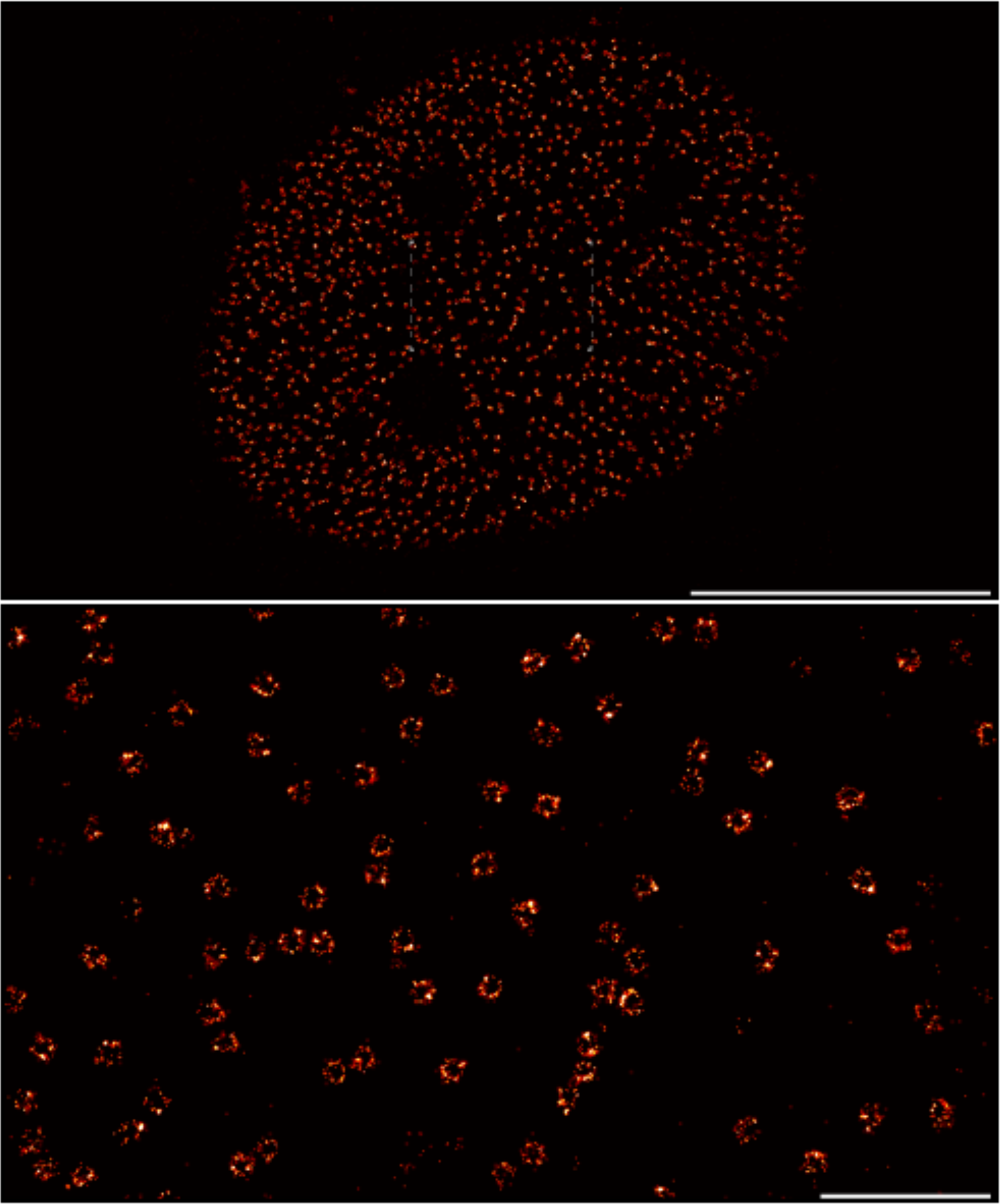
Live-cell SMLM on Nup96-mMaple. Scale bars 10 μm (upper panel) and 1 μm (lower panel).

**Supplementary Figure 5:**
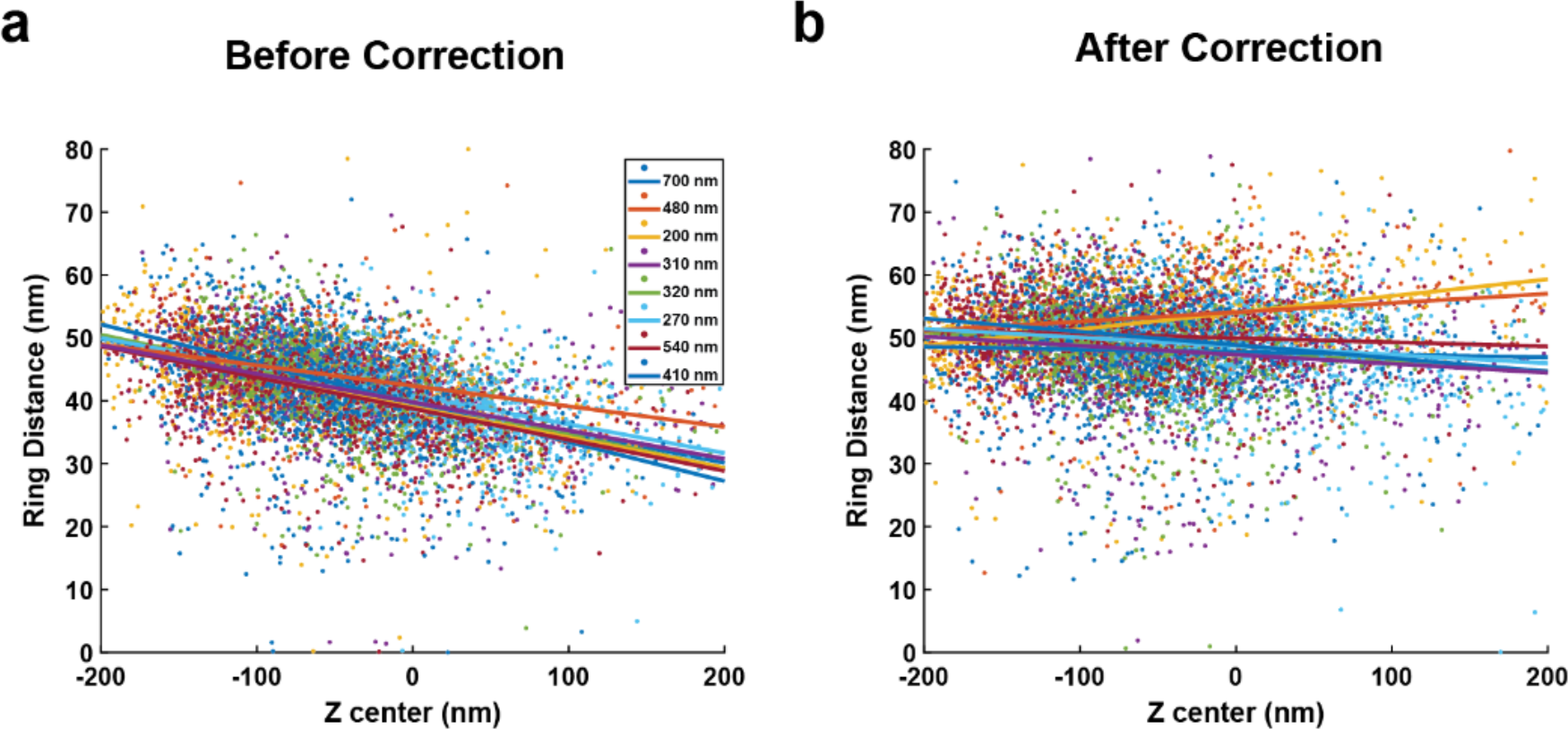
Depth induced aberrations lead to local deformations in z (related to Figure 2j). Distance between rings plotted vs the z-position a) before correction and b) after correction. 7234 NPCs from 8 cells at different depths are shown. Straight lines are the linear fit of the ring distance for each cell. Before correction these values are highly correlated (Pearson coefficient −0.27 ± 0.10), after correcting for the localization errors, the correlation is reduced (Pearson coefficient 0.04 ± 0.15).

**Supplementary Figure 6:**
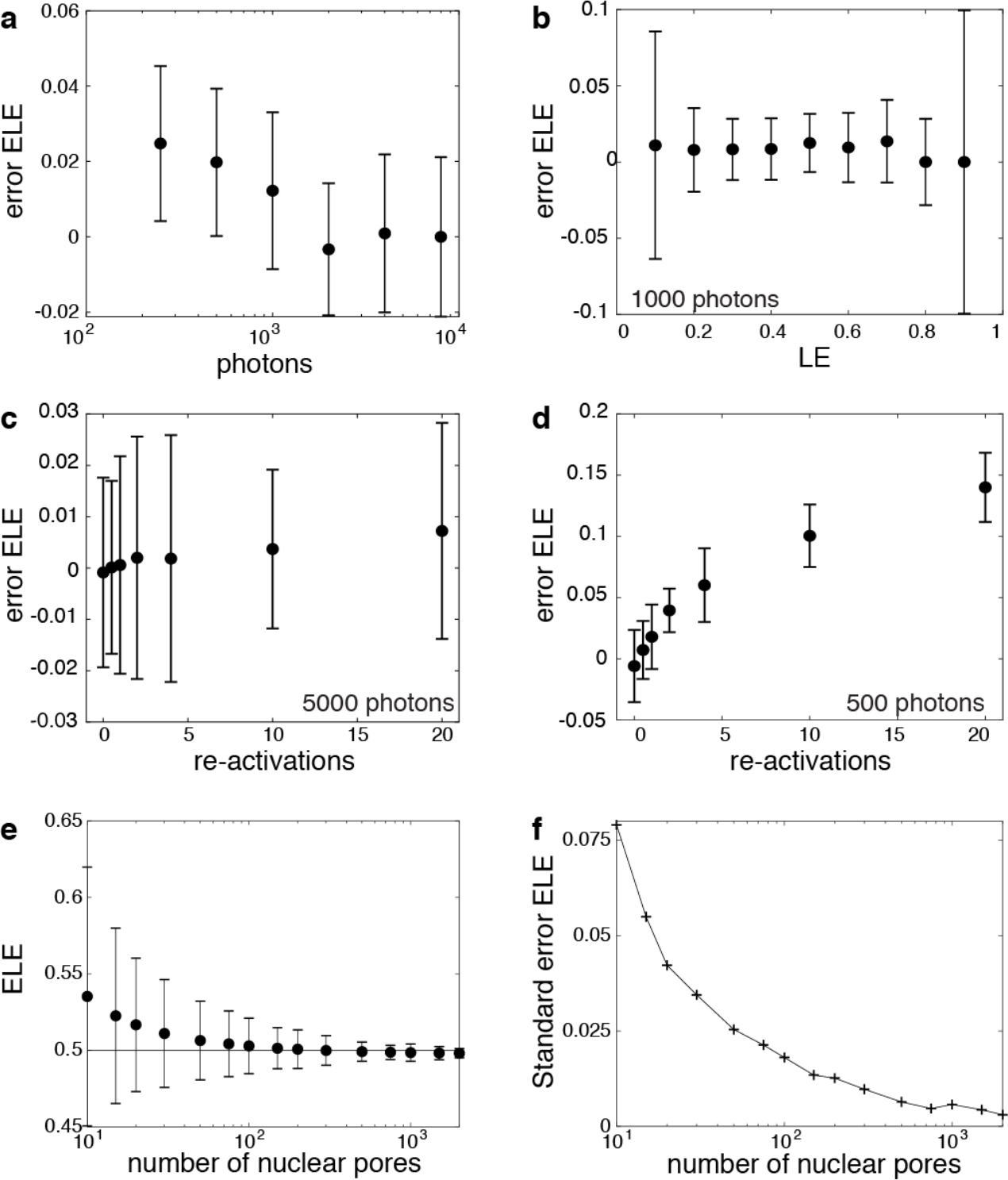
Simulations for determining the ELE. Error in determining ELE (inferred ELE – true ELE) in dependence on (**a**) the brightness of the fluorophores, (**b**) the labeling efficiency, (**c**) the number of re-activations for bright (5000 photons) fluorophores and for (**d**) dim (500 photons) fluorophores and (**e**) in dependence on the number of nuclear pores analyzed. (**f**) statistical accuracy (SEM) in determining the ELE in dependence on the number of nuclear pores. Unless otherwise indicated the simulation parameters were: labeling efficiency 0.5 (**a,e,f**) and 0.3 (**c,d**), number of photons = 5000, on average 1 re-activation, background 20 photons, 900 nuclear pores. Error bars denote mean ± SD.

**Supplementary Figure 7:**
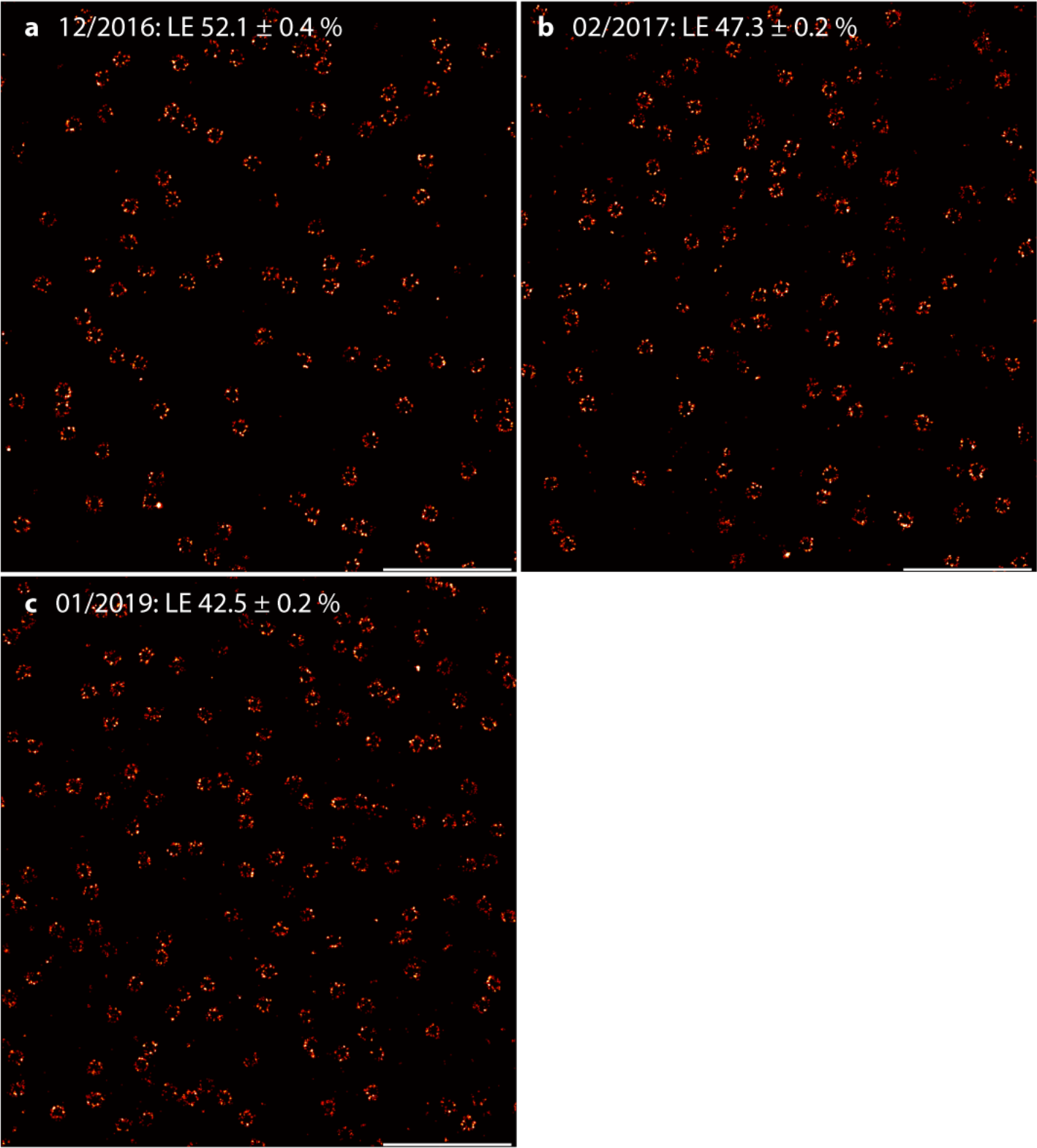
Fixed and labeled samples are stable. ELE of Nup107-SNAP labeled with BG-AF647 and stored at 4°C imaged (**a**) **on the day** of sample preparation, (**b**) **two months** after sample preparation and (**c**) **two years** after sample preparation. Scale bars 1 μm.

**Supplementary Figure 8:**
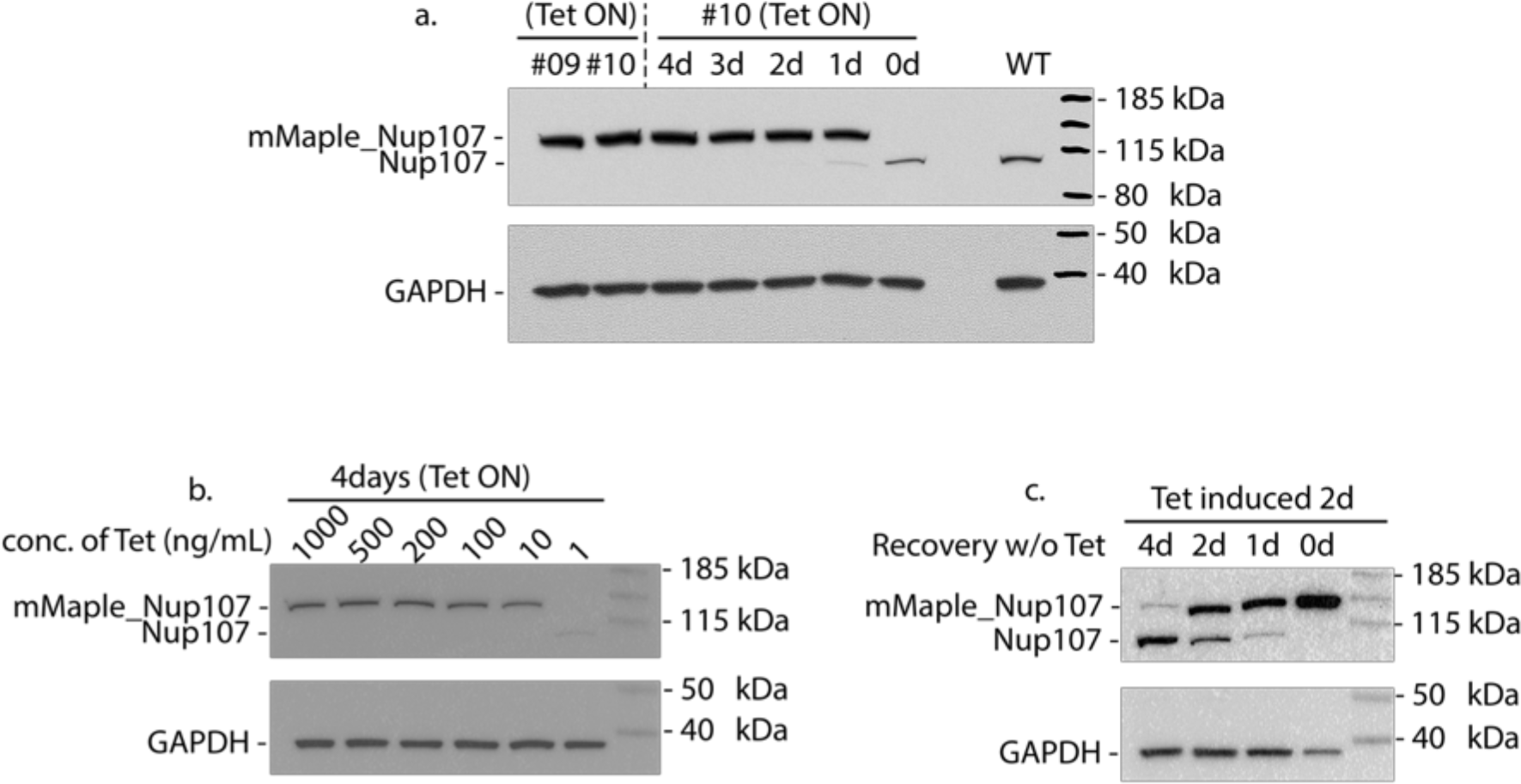
Characterization of stable knock-in HEK293T mMaple-Nup107 cell line. (**a**) **Lanes 1 & 2:** 2 different cell line clones under tetracycline (tet) induction. **Lanes 3-7:** clone no. 10 under tet induction for 4-0 days. **Lane 8:** blank. **Lane 9:** HEK293T wildtype. **Lane 10:** ladder. GAPDH is used as loading control. Clone 10 was used for all subsequent experiments and 2 days of tet induction was sufficient to knock down endogenous Nup107 and induce expressions of mMaple-Nup107. (**b**) Dose titration of tet concentration to induce mMaple-Nup107 expression. 10 ng/mL was sufficient to knock down endogenous Nup107 and induce expressions of mMaple-Nup107. Increasing tet concentration resulted in increasing expression of mMaple-Nup107. A tet concentration of 1 mg/mL was used for all subsequent experiments. **c:** Recovery of endogenous expression of Nup107 after removal of tet from media. Expression of endogenous Nup107 can be observed after 1 day off tet induction with an increasing trend the longer tet is removed. The opposite trend is observed for expression levels of mMaple-Nup107.

## References

1. Hess, S. T., Girirajan, T. P. K. & Mason, M. D. Ultra-High Resolution Imaging by Fluorescence Photoactivation Localization Microscopy. Biophys. J. 91, 4258–4272 (2006).

2. Betzig, E. et al. Imaging Intracellular Fluorescent Proteins at Nanometer Resolution. Science 313, 1642–1645 (2006).

3. Rust, M. J., Bates, M. & Zhuang, X. Sub-diffraction-limit imaging by stochastic optical reconstruction microscopy (STORM). Nat. Methods 3, 793–795 (2006).

4. Xu, K., Zhong, G. & Zhuang, X. Actin, spectrin, and associated proteins form a periodic cytoskeletal structure in axons. Science 339, 452–456 (2013).

5. Mund, M. et al. Systematic Nanoscale Analysis of Endocytosis Links Efficient Vesicle Formation to Patterned Actin Nucleation. Cell 174, 884–896.e17 (2018).

6. Szymborska, A. et al. Nuclear Pore Scaffold Structure Analyzed by Super-Resolution Microscopy and Particle Averaging. Science 341, 655–658 (2013).

7. Ball, G. et al. SIMcheck: a Toolbox for Successful Super-resolution Structured Illumination Microscopy. Sci. Rep. 5, 15915 (2015).

8. Culley, S. et al. Quantitative mapping and minimization of super-resolution optical imaging artifacts. Nat. Methods 15, 263–266 (2018).

9. Sage, D. et al. Super-resolution fight club: A broad assessment of 2D & 3D single-molecule localization microscopy software: bioRxiv 362517 (2018). doi:10.1101/362517

10. Steinhauer, C., Jungmann, R., Sobey, T. L., Simmel, F. C. & Tinnefeld, P. DNA Origami as a Nanoscopic Ruler for Super-Resolution Microscopy. Angew. Chem. Int. Ed. 48, 8870–8873 (2009).

11. Iinuma, R. et al. Polyhedra self-assembled from DNA tripods and characterized with 3D DNA-PAINT. Science 344, 65–69 (2014).

12. van de Linde, S. et al. Direct stochastic optical reconstruction microscopy with standard fluorescent probes. Nat. Protoc. 6, 991–1009 (2011).

13. Huang, B., Jones, S. A., Brandenburg, B. & Zhuang, X. Whole-cell 3D STORM reveals interactions between cellular structures with nanometer-scale resolution. Nat. Methods 5, 1047–1052 (2008).

14. Huang, B., Wang, W., Bates, M. & Zhuang, X. Three-dimensional super-resolution imaging by stochastic optical reconstruction microscopy. Science 319, 810–813 (2008).

15. Pengo, T., Olivier, N. & Manley, S. Away from resolution, assessing the information content of super-resolution images. ArXiv150105807 Phys. (2015).

16. von Appen, A. et al. In situ structural analysis of the human nuclear pore complex. Nature 526, 140–143 (2015).

17. Löschberger, A. et al. Super-resolution imaging visualizes the eightfold symmetry of gp210 proteins around the nuclear pore complex and resolves the central channel with nanometer resolution. J. Cell Sci. 125, 570–575 (2012).

18. Löschberger, A., Franke, C., Krohne, G., van de Linde, S. & Sauer, M. Correlative super-resolution fluorescence and electron microscopy of the nuclear pore complex with molecular resolution. J. Cell Sci. 127, 4351–4355 (2014).

19. Koch, B. et al. Generation and validation of homozygous fluorescent knock-in cells using CRISPR–Cas9 genome editing. Nat. Protoc. 13, 1465–1487 (2018).

20. McEvoy, A. L. et al. mMaple: A Photoconvertible Fluorescent Protein for Use in Multiple Imaging Modalities. PLoS ONE 7, e51314 (2012).

21. Keppler, A., Pick, H., Arrivoli, C., Vogel, H. & Johnsson, K. Labeling of fusion proteins with synthetic fluorophores in live cells. Proc. Natl. Acad. Sci. U. S. A. 101, 9955–9959 (2004).

22. Los, G. V. et al. HaloTag: a novel protein labeling technology for cell imaging and protein analysis. ACS Chem. Biol. 3, 373–382 (2008).

23. Klar, T. A., Jakobs, S., Dyba, M., Egner, A. & Hell, S. W. Fluorescence microscopy with diffraction resolution barrier broken by stimulated emission. Proc. Natl. Acad. Sci. U. S. A. 97, 8206–8210 (2000).

24. Chen, F., Tillberg, P. W. & Boyden, E. S. Expansion microscopy. Science 347, 543–548 (2015).

25. Gustafsson, M. G. L. Surpassing the lateral resolution limit by a factor of two using structured illumination microscopy. J. Microsc. 198, 82–87 (2000).

26. Gustafsson, N. et al. Fast live-cell conventional fluorophore nanoscopy with ImageJ through super-resolution radial fluctuations. Nat. Commun. 7, 12471 (2016).

27. Pesce, L., Cozzolino, M., Lanzanò, L., Diaspro, A. & Bianchini, P. Enigma at the nanoscale: can the NPC act as an intrinsic reporter for isotropic expansion microscopy? bioRxiv (2018). doi:10.1101/449702

28. Heilemann, M. et al. Subdiffraction-resolution fluorescence imaging with conventional fluorescent probes. Angew. Chem. Int. Ed Engl. 47, 6172–6176 (2008).

29. Demmerle, J., Wegel, E., Schermelleh, L. & Dobbie, I. M. Assessing resolution in super-resolution imaging. Methods 88, 3–10 (2015).

30. Nieuwenhuizen, R. P. J. et al. Measuring image resolution in optical nanoscopy. Nat. Methods 10, 557–562 (2013).

31. Li, Y., Wu, Y.-L., Hoess, P., Mund, M. & Ries, J. Depth-dependent PSF calibration and aberration correction for 3D single-molecule localization. bioRxiv 555730 (2019). doi:10.1101/555730

32. Durisic, N., Laparra-Cuervo, L., Sandoval Álvarez, Á., Borbely, J. S. & Lakadamyali, M. Single-molecule evaluation of fluorescent protein photoactivation efficiency using an in vivo nanotemplate. Nat. Methods 11, 156–162 (2014).

33. Fricke, F., Beaudouin, J., Eils, R. & Heilemann, M. One, two or three? Probing the stoichiometry of membrane proteins by single-molecule localization microscopy. Sci. Rep. 5, 14072 (2015).

34. Zanacchi, F. C. et al. A DNA origami platform for quantifying protein copy number in super-resolution. Nat. Methods 14, 789–792 (2017).

35. Ries, J., Kaplan, C., Platonova, E., Eghlidi, H. & Ewers, H. A simple, versatile method for GFP-based super-resolution microscopy via nanobodies. Nat. Methods 9, 582–584 (2012).

36. Grimm, J. B. et al. Bright photoactivatable fluorophores for single-molecule imaging. Nat. Methods 13, 985–988 (2016).

37. Klehs, K. et al. Increasing the Brightness of Cyanine Fluorophores for Single-Molecule and Superresolution Imaging. ChemPhysChem 15, 637–641 (2014).

38. Dempsey, G. T., Vaughan, J. C., Chen, K. H., Bates, M. & Zhuang, X. Evaluation of fluorophores for optimal performance in localization-based super-resolution imaging. Nat. Methods 8, 1027–1036 (2011).

39. Hartwich, T. M. et al. A stable, high refractive index, switching buffer for super-resolution imaging. bioRxiv (2018). doi:10.1101/465492

40. Ori, A. et al. Cell type-specific nuclear pores: a case in point for context-dependent stoichiometry of molecular machines. Mol. Syst. Biol. 9, 648–648 (2013).

41. Otsuka, S. et al. Nuclear pore assembly proceeds by an inside-out extrusion of the nuclear envelope. eLife 5, (2016).

42. Annibale, P., Vanni, S., Scarselli, M., Rothlisberger, U. & Radenovic, A. Quantitative photo activated localization microscopy: unraveling the effects of photoblinking. PLoS ONE 6, e22678 (2011).

43. Lee, S.-H., Shin, J. Y., Lee, A. & Bustamante, C. Counting single photoactivatable fluorescent molecules by photoactivated localization microscopy (PALM). pnas.org

44. Rollins, G. C., Shin, J. Y., Bustamante, C. & Pressé, S. Stochastic approach to the molecular counting problem in superresolution microscopy. Proc. Natl. Acad. Sci. U. S. A. 112, E110–E118 (2014).

45. Finan, K., Raulf, A. & Heilemann, M. A set of homo-oligomeric standards allows accurate protein counting. Angew. Chem. Int. Ed Engl. 54, 12049–12052 (2015).

46. Puchner, E. M., Walter, J. M., Kasper, R., Huang, B. & Lim, W. A. Counting molecules in single organelles with superresolution microscopy allows tracking of the endosome maturation trajectory. Proc. Natl. Acad. Sci. U. S. A. 110, 16015–16020 (2013).

47. Kim, S. J. et al. Integrative structure and functional anatomy of a nuclear pore complex. Nature 555, 475–482 (2018).

48. Rajoo, S., Vallotton, P., Onischenko, E. & Weis, K. Stoichiometry and compositional plasticity of the yeast nuclear pore complex revealed by quantitative fluorescence microscopy. Proc. Natl. Acad. Sci. U. S. A. 115, E3969–E3977 (2018).

49. Wang, S., Moffitt, J. R., Dempsey, G. T., Xie, X. S. & Zhuang, X. Characterization and development of photoactivatable fluorescent proteins for single-molecule-based superresolution imaging. Proc. Natl. Acad. Sci. U. S. A. 111, 8452–8457 (2014).

50. Beck, M., Lučić, V., Förster, F., Baumeister, W. & Medalia, O. Snapshots of nuclear pore complexes in action captured by cryo-electron tomography. Nature 449, 611–615 (2007).

51. Deschamps, J., Mund, M. & Ries, J. 3D superresolution microscopy by supercritical angle detection. Opt. Express 22, 29081–29091 (2014).

52. Gautier, A., Johnsson, K. & O’Hare, H. AGT/SNAP-Tag: A Versatile Tag for Covalent Protein Labeling. in Probes and Tags to Study Biomolecular Function (ed. Miller, L. W.) 89–107 (Wiley-VCH Verlag GmbH & Co. KGaA, 2008). doi:10.1002/9783527623099.ch5

53. Erdmann, R. S. et al. Labeling Strategies Matter for Super-Resolution Microscopy: A Comparison between HaloTags and SNAP-tags. Cell Chem. Biol. (2019). doi:10.1016/j.chembiol.2019.01.003

54. Sun, X. et al. Development of SNAP-Tag Fluorogenic Probes for Wash-Free Fluorescence Imaging. ChemBioChem 12, 2217–2226 (2011).

55. Pleiner, T. et al. Nanobodies: site-specific labeling for super-resolution imaging, rapid epitope-mapping and native protein complex isolation. eLife 4, e11349 (2015).

56. Göttfert, F. et al. Strong signal increase in STED fluorescence microscopy by imaging regions of subdiffraction extent. Proc. Natl. Acad. Sci. U. S. A. 114, 2125–2130 (2017).

57. Tillberg, P. W. et al. Protein-retention expansion microscopy of cells and tissues labeled using standard fluorescent proteins and antibodies. Nat. Biotechnol. 34, 987–992 (2016).

58. Mund, M., Kaplan, C. & Ries, J. Localization microscopy in yeast. Methods Cell Biol. 123, 253–271 (2014).

59. Khmelinskii, A., Meurer, M., Duishoev, N., Delhomme, N. & Knop, M. Seamless Gene Tagging by Endonuclease-Driven Homologous Recombination. PLoS ONE 6, e23794 (2011).

60. Deschamps, J., Rowald, A. & Ries, J. Efficient homogeneous illumination and optical sectioning for quantitative single-molecule localization microscopy. Opt. Express 24, 28080–28090 (2016).

61. Ong, W. Q., Citron, Y. R., Schnitzbauer, J., Kamiyama, D. & Huang, B. Heavy water: a simple solution to increasing the brightness of fluorescent proteins in super-resolution imaging. Chem. Commun. Camb. Engl. 51, 13451–13453 (2015).

62. Heilemann, M., Margeat, E., Kasper, R., Sauer, M. & Tinnefeld, P. Carbocyanine dyes as efficient reversible single-molecule optical switch. J. Am. Chem. Soc. 127, 3801–3806 (2005).

63. Bates, M., Blosser, T. & Zhuang, X. Short-Range Spectroscopic Ruler Based on a Single-Molecule Optical Switch. Phys Rev Lett 94, 108101 (2005).

64. Li, Y. et al. Real-time 3D single-molecule localization using experimental point spread functions. Nat. Methods 15, 367–369 (2018).

65. Mortensen, K. I., Churchman, L. S., Spudich, J. A. & Flyvbjerg, H. Optimized localization analysis for single-molecule tracking and super-resolution microscopy. Nat. Methods 7, 377–381 (2010).

